# Omics-informed CNV calls reduce false positive rate and improve power for CNV-trait associations

**DOI:** 10.1101/2022.02.07.479374

**Authors:** Maarja Lepamets, Chiara Auwerx, Margit Nõukas, Annique Claringbould, Eleonora Porcu, Mart Kals, Tuuli Jürgenson, Estonian Biobank Research Team, Andrew Paul Morris, Urmo Võsa, Murielle Bochud, Silvia Stringhini, Cisca Wijmenga, Lude Franke, Hedi Peterson, Jaak Vilo, Kaido Lepik, Reedik Mägi, Zoltán Kutalik

**Affiliations:** Estonian Genome Centre, Institute of Genomics, University of Tartu, Tartu, Estonia; Institute of Molecular and Cell Biology, University of Tartu, Tartu, Estonia; Center for Integrative Genomics, University of Lausanne, Lausanne, Switzerland; Department of Computational Biology, University of Lausanne, Lausanne, Switzerland; Swiss Institute of Bioinformatics, Lausanne, Switzerland; Center for Primary Care and Public Health (Unisanté), Department of epidemiology and health systems, University of Lausanne, Lausanne, Switzerland; Structural and Computational Biology Unit, EMBL, Heidelberg, Germany; Institute for Molecular Medicine Finland, FIMM, HiLIFE, University of Helsinki, Helsinki, Finland; Institute of Mathematics and Statistics, University of Tartu, Tartu, Estonia; Centre for Genetics and Genomics Versus Arthritis, Centre for Musculoskeletal Research, Division of Musculoskeletal and Dermatological Sciences, The University of Manchester, Manchester, UK; Unit of Population Epidemiology, Division of Primary Care, Geneva, Switzerland; University of Groningen, University Medical Center Groningen, Department of Genetics, Groningen, The Netherlands; Oncode Institute, Utrecht, The Netherlands; Institute of Computer Science, University of Tartu, Tartu, Estonia

**Keywords:** anthropometric traits, copy number variation, gene expression, methylation, multi-omics, PennCNV, structural variation, whole genome sequencing

## Abstract

Copy number variations (CNV) are believed to play an important role in a wide range of complex traits but discovering such associations remains challenging. Whilst whole genome sequencing (WGS) is the gold standard approach for CNV detection, there are several orders of magnitude more samples with available genotyping microarray data. Such array data can be exploited for CNV detection using dedicated software (e.g., PennCNV), however these calls suffer from elevated false positive and negative rates. In this study, we developed a CNV quality score that weights PennCNV calls (pCNV) based on their likelihood of being true positive. First, we established a measure of pCNV reliability by leveraging evidence from multiple omics data (WGS, transcriptomics and methylomics) obtained from the same samples. Next, we built a predictor of omics-confirmed pCNVs, termed *omics-informed quality score* (OQS), using only PennCNV software output parameters. Promisingly, OQS assigned to pCNVs detected in close family members was up to 35% higher than the OQS of pCNVs not carried by other relatives (P < 3.0×10^−90^), outperforming other scores. Finally, in an association study of four anthropometric traits in 89,516 Estonian Biobank samples, the use of OQS led to a relative increase in the trait variance explained by CNVs of up to 34% compared to raw pCNVs or previous quality scores. Overall, we put forward a flexible framework to improve any CNV detection method leveraging multi-omics evidence, applied it to improve PennCNV calls and demonstrated its utility by improving the statistical power for downstream association analyses.

## Introduction

Copy number variations (CNV) are unbalanced structural variations that alter the dosage of genomic regions via deletion and duplication events. Approximately 9.5% of the human genome is subject to CNVs^1^, which vary in length, ranging from a few dozens to several millions of base pairs (bp) in length. CNVs tend to have more severe phenotypic consequences compared to single nucleotide variations (SNV) as, due to their larger size, they can encompass entire coding regions.

CNVs have been associated with a number of conditions including autism^2^, schizophrenia^3^, neurodegenerative disorders^4,5^, and cancer^6^. A number of large recurrent deletions and duplications have been combined into the DECIPHER CNV syndromes list^7^. Importantly, incomplete penetrance of several syndromic CNVs has been established by studying large population biobanks, where CNV load was shown to increase the risk of obesity, physical or cognitive impairment, and congenital malformations while lowering educational attainment and socio-economic status^8–11^. In parallel, CNV genome-wide association studies (GWAS) have been conducted on numerous diagnoses^12,13^ and medically relevant continuous traits^11,14,15^, including a large meta-analysis on anthropometric measurements^16^, revealing the important role of CNVs in shaping the human phenome.

Over the years, multiple CNV detection algorithms have been developed for CNV detection from SNV genotyping microarray probe intensities^17^. Currently, PennCNV^18^ is the most widely used software for genotyping array-based calling. For each sample, a Hidden Markov Model (HMM)-based algorithm uses overall signal intensity and continuous allelic intensity at polymorphic probes to estimate the probability of a hidden copy number state at this genomic location. Unfortunately, CNV regions found by different array-based detection methods only agree in about 20% of cases^19^, indicating the high likelihood of false positive calls. To counter this, various filtering strategies have been employed, usually by setting cut-off values to combinations of parameters including number of CNVs per sample, minimum CNV length, probe-density and PennCNV confidence score^9,12,18,20,21^. Filtering based on arbitrary thresholds is suboptimal and a continuous CNV quality score that predicts the probability that a CNV region is a consensus call between PennCNV, QuantiSNP^22^ and CNVpartition (an Illumina developed GenomeStudio software plug-in, https://support.illumina.com/downloads/genomestudio-2-0-plug-ins.html) has been proposed^19^. We refer to this as consensus-based quality score (cQS).

Still, cQS relies only on a single input dataset (i.e., microarray data). An alternative strategy to improve CNV calling can be to incorporate various types of omics datasets. Previously, software have been developed to infer CNVs from high-density DNA methylation arrays^23,24^ or RNA sequencing data of highly and stably expressed genes^25^. While promising, none of these approaches were developed with the intent of performing scalable and reliable genome-wide CNV detection in large biobanks.

To fill this gap, we propose a method to improve the detection of false positive CNV calls amongst PennCNV output by discriminating between high quality (true) and low quality (false) CNV regions based on multi-omics data (**Figure 1A**). Specifically, we checked if PennCNV calls (pCNV) (1) are detectable by WGS, (2) alter the expression levels of overlapping genes in the expected direction (i.e., decreased by deletions, increased by duplications) and/or (3) alter the total methylation probe intensity of overlapping CpG sites in the expected direction. We built a predictor of CNV quality inferred from WGS, transcriptomics and methylomics, solely based on PennCNV software output parameters in these samples assayed by multiple omics technologies. Predicted omics-informed quality scores (OQS) distinguish high from low quality CNVs even in samples for which only SNV genotyping microarray data are available. We show that OQS reduces false discovery rate and improves CNV-trait association discovery compared to both raw PennCNV calls and cQS^19^ in regions with variable CNV quality.

**Figure 1.**
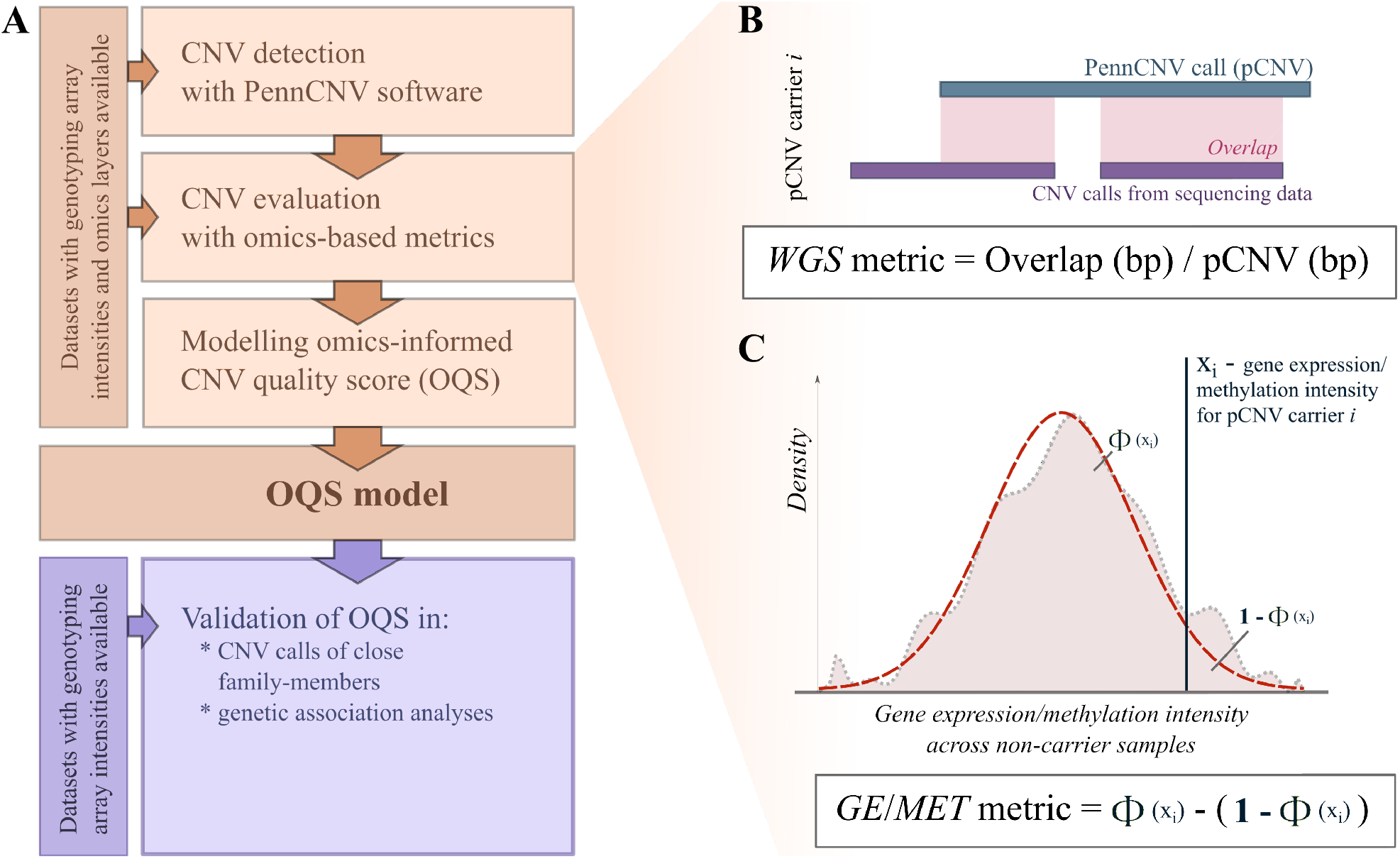
(A) Quality estimation and modelling pipeline for PennCNV copy number variation calls (pCNV). The pCNV quality metrics are estimated based on (B) whole genome sequencing (WGS) data and (C) gene expression (GE) and/or overall methylation intensity (MET) of genes/CpG sites overlapping the corresponding CNV call. (B) WGS metric is a fraction of pCNV that can be mapped to WGS CNVs of the same individual. (C) To calculate GE/MET metrics, the reference distribution of expression/intensity based on non-carriers (pink area) is approximated to standard normal distribution (red dashed line) and the Z score of the expression/intensity of each pCNV carrier (x_i_) is compared to it one at a time. The metric is a difference between the fraction of non-carriers with the corresponding value ≤x_i_ and those with the corresponding value >x_i_ and captures how extreme x_i_ is compared to the reference distribution of non-carriers. In case a pCNV overlaps with several genes/CpG sites, the metric values are averaged over them.

## Methods

### Cohorts

*Estonian Biobank* (EstBB; data freeze 2021/01/08; **Supplementary Note S1, Table 1**) is an Estonian population-based cohort that consists of ∼200,000 adults (≥18 years of age at recruitment)^26^. About 7,750 individuals are genotyped on Illumina Infinium OmniExpress-24 genotyping array. A subset of these samples (referred to as EstBB-MO) has one or more of the following omics datasets available: 30x coverage whole-genome sequencing (WGS), RNA sequencing^27^ and/or methylation data (Illumina Infinium Human Methylation 450k Beadchip).

**Table 1.**
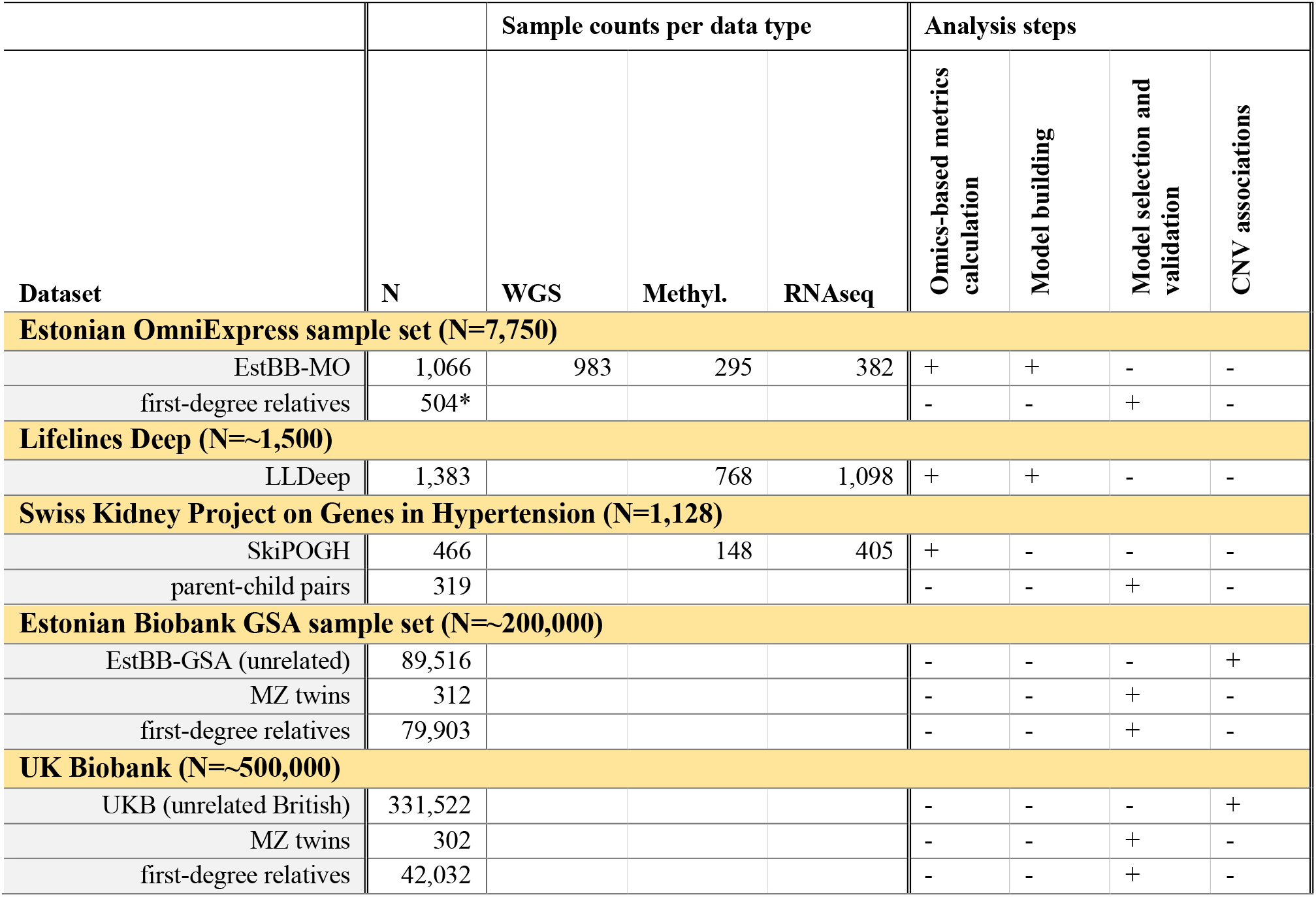
Overview of datasets and final sample sizes used in the analyses. *Estonian OmniExpress first-degree relatives do not overlap with EstBB-MO samples.

Additionally, the full EstBB cohort is genotyped on Illumina Global Screening Array (GSA). All participants signed a broad informed consent and analyses were carried out under ethical approval 1.1-12/624 from the Estonian Committee on Bioethics and Human Research and data release N05 from the EstBB.

#### LifeLines Deep

(LLDeep; **Supplementary Note S2, Table 1**) is a deeply phenotyped ∼1,500 individual subset of the Dutch population cohort LifeLines^28^. LLDeep samples are genotyped on HumanCytoSNP-12 array and the majority of them have either RNA sequencing^29^ or methylation data (Illumina Infinium Human Methylation 450k Beadchip)^30^ available. The LifeLines DEEP study was approved by the ethics committee of the University Medical Centre Groningen. All participants provided a written informed consent.

#### Swiss Kidney Project on Genes in Hypertension

(SkiPOGH; **Supplementary Note S3, Table 1**) is a Swiss family and population-based cohort of 1,128 individuals from 273 families recruited to study the genetic determinants of blood pressure^31^. The samples were genotyped on Illumina 2.5 array. RNA sequencing and methylation array (Illumina Infinium Human Methylation 450k) data were available for a subset of participants^32^. The study was approved by the competent institutional ethics committees in Bern, Geneva and Lausanne. All participants signed a written informed consent.

#### UK Biobank

(UKB; phenotype data freeze 2018/03/22; **Supplementary Note S4, Table 1**) is a cohort of ∼500,000 individuals from the United Kingdom^33^. The majority of samples (∼450,000) are genotyped on Affymetrix UK Biobank Axiom array while the rest (∼50,000) are genotyped on Affymetrix UK BiLEVE Axiom array. Participants signed a broad informed consent, and the data are accessed through application numbers 17085 and 16389.

### Data preparation

#### Sample sets

We included three independent datasets – LLDeep, SkiPOGH and a subset of Estonian samples (EstBB-MO) – in CNV quality calculations and modelling. Each of these datasets had additional omics data (WGS, methylation arrays and/or RNA sequencing) available. For model selection and validation steps we extracted monozygotic (MZ) twins and first-degree relatives from the EstBB and the UKB, and parent-child pairs from the SkiPOGH. Finally, we extracted unrelated quality-controlled EstBB-GSA and UKB samples for CNV association analyses. Datasets and their usage for various analyses are summarised in **Table 1**. Sample quality control steps are summarised in **Supplementary Notes S1-S4**.

#### CNV detection

We used PennCNV^18^ as our main CNV detection algorithm due to its popularity (PubMed citations: PennCNV: 885; QuantiSNP^22^: 279; Birdsuite^34^: 466; 2021/09/08). We detected putative autosomal CNV regions (pCNV) for EstBB, LLDeep, SkiPOGH, and UKB datasets as previously described^19^ (**Supplementary Notes S1-S4**). For each sample, we obtained the pCNV together with the values of four CNV-specific and nine sample-specific parameters described in **Table S1**. In all datasets, we filtered out samples with more than 200 pCNVs and pCNVs larger than 10Mbp, as these are likely to be either samples with poor genotyping quality or extreme cases that might distort the analysis. Additionally, we detected CNVs from EstBB WGS reads (WGS-CNVs) using the Genome STRiP^35^ discovery pipeline (version 2.00.1611; **Supplementary Note S5**). All genomic coordinates are in GRCh37 build version.

#### Methylation and RNA-sequencing data pre-processing

We obtained methylation intensities (Infinium Human Methylation 450k Beadchip) and RNA-sequencing data for EstBB-MO, LLDeep and SkiPOGH datasets. The data preparation is in detail described in **Supplementary Notes S6-S7**. Briefly, after quality control step we corrected for age, sex, batch, blood cell counts and genotype principal components (PCs) where applicable. Additionally, we corrected for PCs calculated based on methylation/gene expression data (**Figures S1 and S2**). Gene expression residuals were further corrected for independent expression quantitative trait loci (eQTL) within 500kbp of the gene.

### CNV quality metrics based on multi-omics data

#### WGS quality metric

WGS data were available for a subset of EstBB-MO samples (N=979). For each pCNV in these individuals, we defined a *WGS* metric as the fraction of the pCNV (in bps) overlapping with WGS-CNVs in the same sample (**Figure 1B**). Metric calculation was restricted to pCNV deletions and duplications longer than 1kb and 2kb, respectively, as we did not detect shorter WGS-CNVs (**Supplementary Note S5**).

#### MET quality metric

Methylation intensity data were used to assess CNV quality in EstBB-MO, LLDeep and SkiPOGH datasets. For each methylation probe passing the pre-processing steps (**Supplementary Note S6**), we used the samples with no pCNVs overlapping the corresponding CpG site (i.e., non-carriers) to construct the approximately Gaussian reference distribution of site overall intensity (sum of the methylated and un-methylated intensities). For each carrier, we then transformed its CpG site overall intensity into a Z score by using the mean and standard deviation of the constructed reference distribution. We denoted the quality metric based on the methylation data for the *i*-th pCNV (*pCNV*_*i*_) across all its overlapping CpG sites as *MET*_*i*_ ∈ [−1; 1] and calculated its value as:

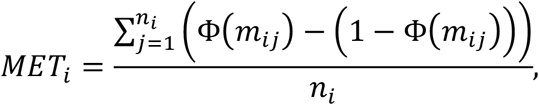

where *n*_*i*_ is the total number of CpG sites overlapping *pCNV*_*i*_, Φ is the cumulative distribution function of the standard normal distribution, and *m*_*ij*_ is the Z score calculated for the *j*-th CpG site overlapping *pCNV*_*i*_ (**Figure 1C**). The proposed measure captures how extreme an observed methylation intensity is compared to that of the bulk of the samples (assumed to be copy-neutral), equivalent to a signed 2-sided tail-probability. We expected *MET*_*i*_ <0for deletions and *MET*_*i*_ *>* 0 for duplications. If this was not the case, *MET*_*i*_ was set to zero. Finally, *MET*_*i*_ was converted to its absolute value such that *MET*_*i*_ ∈ [0; 1].

#### GE quality metric

Gene expression levels from RNA sequencing data were used to assess CNV quality in the EstBB-MO, LLDeep and SkiPOGH datasets. We extracted all the genic regions from Ensemble database (GRCh37) using biomaRt^36^. We retained 10,786 genes passing the pre-processing steps (**Supplementary Note S7**) and having expression correlated to the copy number of the gene (Pearson *R >* 0.1) in an independent dataset^25^. Additionally, over 80% of the genic region was required to overlap the pCNV for the gene to be included in the quality calculations of that pCNV. Expression values of genes with *pCNV*_*i*_ overlap below 80% were marked as missing. Analogously to *MET*_*i*_, we constructed the expression reference distribution based on non-carriers and used its mean and standard deviation to calculate an expression Z score for each carrier. We calculated *GE*_*i*_ ∈ [−1; 1], the *pCNV*_*i*_ quality metric based on gene expression across all its overlapping genes (analogously to the metric for methylations), as:

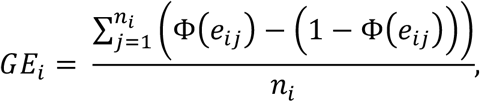

where *n*_*i*_ is the number of genes overlapping at least 80% of the *pCNV*_*i*_ and *e*_*ij*_ is the Z score calculated for the *j*-th gene overlapping *pCNV*_*i*_ (**Figure 1C**). We set to zero *GE*_*i*_ values for deletions with *GE*_*i*_ *>* 0 and duplications with *GE*_*i*_ < 0 and converted all *GE*_*i*_ scores to their absolute values such that *GE*_*i*_ ∈ [0; 1].

#### Combined metric

Let us define the collection of quality metrics ***Q***_***i***_ *=* {*MET*_*i*_, *GE*_*i*_, *WGS*_*i*_}. Further metrics can be defined by their mean, maximum and the measure furthest away from 0.5 (i.e., most extreme; denoted as *EXTR(Q*_*i*_)). We chose *EXTR(Q*_*i*_) as our final combined metric, the motivation being that if one metric clearly indicated the truth status of a pCNV then we would use that metric (even if other metrics are unsure) (**Figure S3**). Note that *EXTR* values are only calculated for pCNVs that have at least two out of three *Q*_*i*_ metrics available.

### CNV quality prediction models

In order to assess CNV quality in samples with no complementary omics data, we fitted prediction models for the values of the previously defined four omics metrics, summarized by **Y**_**i**_ **=** {*MET*_*i*_, *GE*_*i*_, *WGS*_*i*_, *EXTR(Q*_*i*_*)}*.

The set of possible explanatory variables ***X*** = {*X*_1_, *X*_2_, …, *X*_*n*_} included CNV and sample-specific parameters from PennCNV output (CNV length, number of overlapping probes, PennCNV-specific CNV confidence score, number of pCNVs per sample (and its derivations, see **Figure S4**) mean and standard deviation of the allelic intensity ratios and B allele frequencies of a sample, signal waviness factor; **Table S1**) and their interaction terms. We fitted a generalized linear regression model separately on each column of ***Y***:

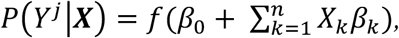

where *Y*^*j*^ is the *j*-th column of **Y** (representing one of the four quality metrics), *β*_0_ is the intercept term and *β*_*k*_ is the effect estimate of explanatory variable *X*_*k*_. We used a quasi-binomial link function *f*, since instead of being strictly binary, our response was a bimodal continuous variable ranging from zero to one.

In order to choose the best subset of ***X***, we implemented a forward stepwise model selection (starting with an empty parameter set) using custom R scripts. Briefly, in each round, a parameter was added if the resulting model minimized the 10-fold cross-validation mean square error (MSE). If adding any of the remaining parameters did not improve the average MSE, the algorithm stopped and returned the existing model. We tested model building with eight different sets of conditions/parameters {*X*^*s*^: *X*^*s*^ ⊂ ***X***} to choose from and repeated the procedure separately for deletions and duplications. The modelling process and the eight parameter sets are characterized in detail in **Supplementary Note S8 and Table S2**.

The model coefficients *β*_*k*_ can be used to predict omics-informed CNV quality scores (OQS) as:

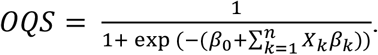

For *X*_*k*_ not included in the final model, *β*_*k*_ is set to be equal to zero.

### Selection of the best omics-informed quality score and comparison to other CNV calls

CNV quality models were fitted as described above for each multi-omics dataset separately (**Table 1**). Altogether, we fitted 24-32 (not necessarily unique) models for both deletions and duplications per dataset (3-4 omics measures x 8 model parameter sets).

To determine the best models, we incorporated family information. We reasoned that the set of ‘familial’ pCNVs present in at least two close family-members (Jaccard index based on overlapping bp count of at least 0.9) contains a higher fraction of true positive CNV calls than the ‘non-familial’ set (no overlap in a relative). Partially overlapping pCNVs with Jaccard index lower than 0.9 were discarded from the calculations. We predicted and compared the OQS values of all familial and non-familial pCNVs from MZ twins from the UKB and EstBB-GSA and parent-child pairs from the SkiPOGH. To select the best models, we maximised the difference of mean OQS values between the two groups, averaged over the three datasets. We validated our best models on first-degree relative pCNVs from Estonian OmniExpress samples, EstBB-GSA and UKB. As a comparison, we calculated these differences for the previously published consensus-based quality score (cQS)^19^.

### CNV association analyses

We compared OQS to raw PennCNV calls and cQS in an association analysis setting by incorporating them into the association models analogously to SNV dosages^16^. We used 89,516 and 331,522 quality-controlled unrelated European individuals from the EstBB-GSA and UKB data, respectively (**Supplementary Note S1 and S4**). We considered 21 CNV-trait pairs (**Table S3**) involving four continuous anthropometric traits – body mass index (BMI), height, weight and waist-to-hip ratio (WHR) – and 13 CNV regions which had previously^16^ shown association P-value <1×10^−4^. Importantly, these associations were obtained using cQS in the first wave of UKB genotype data samples (N=119,873). All phenotypes were inverse normal transformed and corrected for batch, sex, age, age^2^ and PCs prior further analysis (**Supplementary Notes S1 and S4**).

We calculated association Z statistics (estimated effect size over its standard error) and P-values in a probe-by-probe manner across all probes that overlapped with >5 pCNVs inside the 13 regions of interest. The analysis was conducted separately for deletions, duplications and mirroring effects. To model the mirroring effect (i.e., both deletions and duplications have similar effects but in opposite directions), the OQS values for deletions were negated.

Associations were run using linear regression (*lm* function) in custom R scripts. We used a Bonferroni-corrected P-value threshold of 0.05/21=2.38×10^−3^ to determine significance. All regions containing significantly associated probes - except for the 18q21.32 region, which in the EstBB-GSA did not contain the previously reported CNV (**Figure S5**) - were included in the final association comparison step.

Finally, for both datasets (i.e., UKB and EstBB-GSA) and all four phenotypes we estimated the change in explained variance when applying the OQS model, as compared to raw PennCNV values or cQS model. We started by clumping probes originating from significant CNV regions with *R*^2^ *>* 0.5 using *snp_clumping* from the *bigsnpr* R package^37^. Clumping usually prioritizes probes based on association summary statistics or allele frequencies, which in our case are heavily dependent on the applied CNV quality measure. To avoid any bias, we generated a random probe priority order instead. If after clumping we retained *m* probes, we calculated

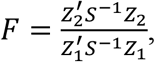

where *Z*_2_ is an array of Z statistics (of clumped probes) from association analysis using OQS, *Z*_1_ is the corresponding array from the comparison analysis (either raw PennCNV or cQS) and *S* is a probe correlation matrix calculated based on raw PennCNV calls. Under the null hypothesis, *F* follows an F distribution with both degrees of freedom equal to *m*. Since *F* depends on the probes retained after the randomised clumping process, we repeated the random clumping 20 times and used the average F-value.

## Results

### Omics-based metrics for CNV quality

We estimated pCNV quality based on methylation (*MET*), gene expression (*GE*) and whole-genome sequencing (*WGS*, only available in EstBB-MO dataset) data, which resulted in up to three independent omics-based CNV quality metrics in three independent datasets (EstBB-MO, LLDeep and SkiPOGH; **Tables 1 and S4**). Within all datasets the metrics were positively correlated with each other (**Figures 2A-B, S6 and S7**). Both *MET* and *GE* metrics had high correlations (Pearson *R* ≥ 0.7) with the *WGS* metric. The correlations between *MET* and *GE* metrics ranged between 0.59 and 0.80 for deletions and 0.33 and 0.57 for duplications. The correlations between all three metrics and previously published consensus-based quality score (cQS)^19^ ranged between 0.17 and 0.55 for deletions and 0.21 and 0.63 for duplications depending on the dataset.

**Figure 2.**
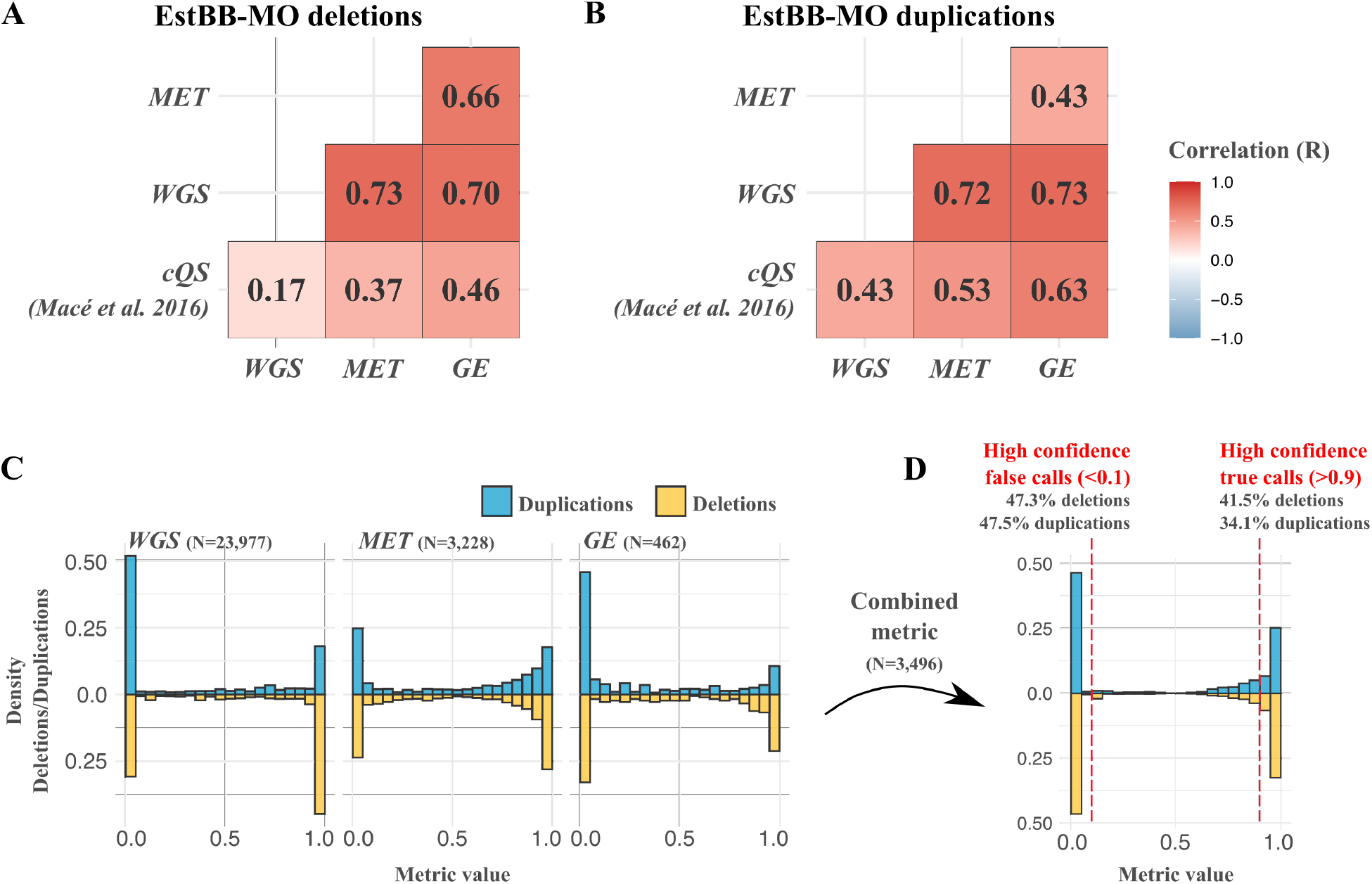
Omics-based metrics – sequencing (WGS), methylation (MET) and gene expression (GE) – and cQS^19^ Pearson correlations for EstBB-MO deletions (A) and duplication (B). Note that the number of pCNVs used in correlation calculations is not identical in each group of metric pairs (**Figure S8**). Bimodal distribution of WGS, MET and GE metrics (C), as well as their combined metric (see Methods) (D) for duplications (blue) and deletions (yellow). The combined metric is calculated for pCNVs that have at least two omics-based metrics available (N=3,496) and the fractions of high confidence false (combined metric <0.1) and true (combined metric >0.9) calls are reported.

All three metrics had bimodal distributions with modes near 0 and 1, which indicates clear differentiation between true and false calls for the majority of pCNVs (**Figures 2C, S6 and S7)**. To retain just one quality metric per pCNV (i.e., combined metric, **Figure 2D**), we retained the metric that was ‘furthest’ from 0.5 (denoted as *EXTR* in **Methods**).

We estimated the precision of PennCNV calls based on the combined metric. In the EstBB-MO dataset, out of 3,496 pCNVs evaluated with the combined metric (1,750 deletions, 1,746 duplications), 47.3% of deletions and 47.5% of duplications had values inferior to 0.1, most likely reflecting false positive calls. In LLDeep and SkiPOGH, the corresponding percentages were 50.5%/28.3% and 70.9%/59.4% for deletions/duplications, respectively (**Table S5**), illustrating the need for CNV quality filtering prior further analyses.

### Prediction models for omics-informed CNV quality scores (OQS)

We built logistic regression models to predict the previously calculated CNV quality metrics based solely on PennCNV output parameters to enable pCNV evaluation in samples lacking multi-omics measurements. Due to a smaller fraction of true positive calls when compared to other datasets, we omitted SkiPOGH from the model-building step but retained it for model validations. Models were evaluated based on their ability to discriminate between pCNVs that were shared (‘familial’, assumed to be true calls) or not (‘non-familial’) between MZ twins in the UKB, EstBB-GSA and parent-child pairs in the SkiPOGH (**Tables S6 and S7**). For both deletions and duplications, the best model was built based on the LLDeep dataset using the combined metric. Models are characterised in **Tables S8 and S9**. We refer to the CNV quality measure (ranging from 0 to 1) predicted by the best models as the omics-informed CNV quality score (OQS).

To validate the OQS, we performed a ‘familial’ *versus* ‘non-familial’ pCNV comparison on first-degree relatives from the Estonian OmniExpress, EstBB-GSA and UKB that did not overlap with individuals used for the CNV quality estimation and model building steps (i.e., samples with other omics data; **Figures 3 and S9**). The average OQS for familial calls ranged between 0.67 and 0.82 for deletions, and between 0.48 and 0.70 for duplications, which was significantly higher (paired Wilcoxon test *P* < 1.4 × 10^−21^) than for cQS (0.27-0.32 in deletions and 0.42-0.53 in duplications). Furthermore, OQS distinguished well between familial and non-familial pCNVs. The difference in OQS between two groups were between 0.22 and 0.35 depending on the dataset (0.16-0.25 for deletions and 0.12-0.48 for duplications; Wilcoxon *P* < 3.0 × 10^−90^). Only in case of EstBB-GSA duplications was the average difference larger with cQS.

**Figure 3.**
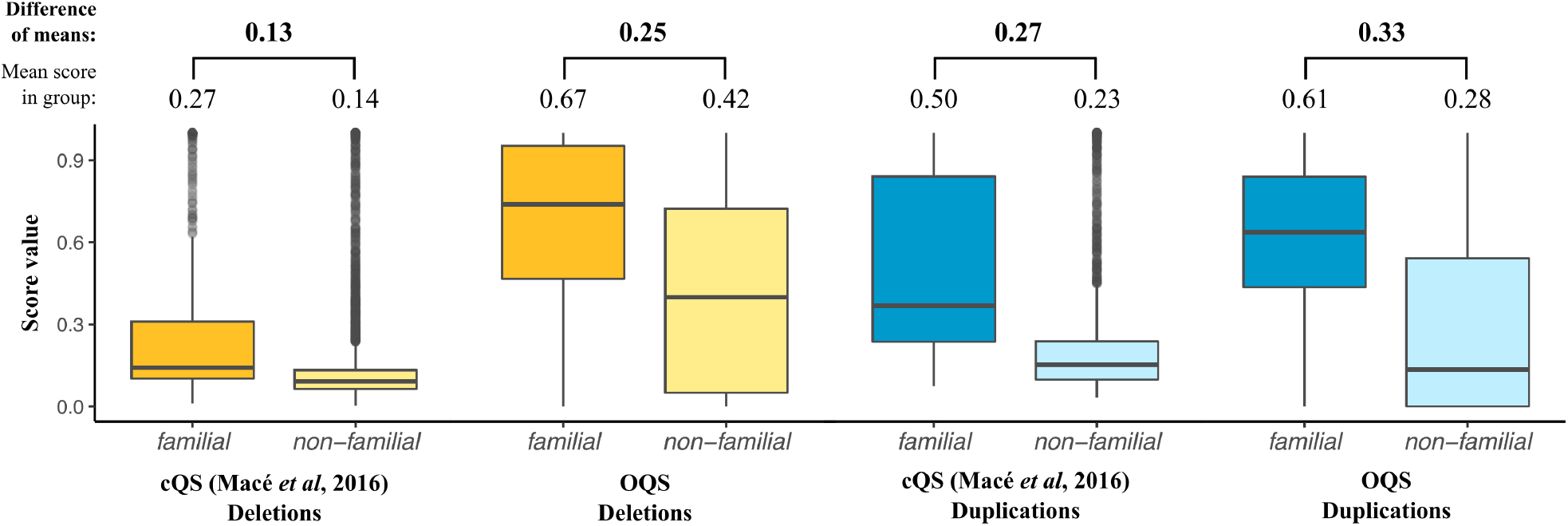
Consensus-based (cQS)^19^ and omics-informed (OQS) CNV quality scores of familial (found in two or more family members) and non-familial deletions (yellow) and duplications (blue) calculated on a subset of Estonian OmniExpress samples (N=504; do not overlap with EstBB-MO). Familial pCNVs are likely true positives while non-familial group contains both true and false positives. The mean score of each pCNV group and their pairwise difference is shown on top of the figure. Compared to cQS the OQS shows higher values for familial pCNVs and larger differences between non-familial and familial pCNV quality. All differences for both scores are significant with P-value < 1×10^−16^ (Wilcoxon test).

As the best models were built on LLDeep dataset, we could use EstBB-MO for out-of-sample validations (**Figure S10**). We found that Pearson correlation coefficients between the combined metric and predicted OQS were 0.70 and 0.57 for deletions and duplication, respectively. The AUC values were 0.91 for deletions and 0.87 for duplications (**Figure S11**).

### Associations between CNV and anthropometric traits

We compared the association results obtained using raw PennCNV calls, cQS^19^ and OQS. Of 21 previously established associations between CNVs and four anthropometric traits (body mass index (BMI), height, weight and waist-to-hip ratio (WHR))^16^, we replicated (P < 2.38×10^−3^) 10 in the EstBB-GSA and 18 in the UKB cohort (**Tables S3, S10 and S11**). For both datasets we calculated the change in variance explained per phenotype when using OQS compared to the other two approaches.

First, we tested mirror type associations where deletions and duplications have similar effects but opposite effect directions. We found that in the EstBB-GSA, OQS led to a relative increase of 2%-34% and 23%-55% in the explained variance compared to raw PennCNV and cQS, respectively, depending on the phenotype (**Figure 4A** and **Table S12**). A good example is an association between the 16p11.2 BP4-BP5 CNV status and BMI (**Figure 4B**), for which alone the relative variance explained increased by 26% and 40% compared to raw PennCNV and cQS, respectively. For deletion-only associations, the relative increase was equally good, up to 33% and 46% compared to raw PennCNV and cQS, respectively (**Figure S12**). For duplication-only analysis, only one BMI-associated region was included and it showed 43% relative increase in explained variance compared to the other approaches. In the UKB, OQS showed improvement compared to raw PennCNV in three out of four phenotypes. However, compared to the cQS the explained variance was decreased in most cases. This was to be expected as the associations incorporated in this study were originally detected using the cQS in a dataset where over 60% of samples were from UKB^16^. None of the changes were statistically significant as the number of independent CNV regions per phenotype was very low, ranging from one to seven.

**Figure 4.**
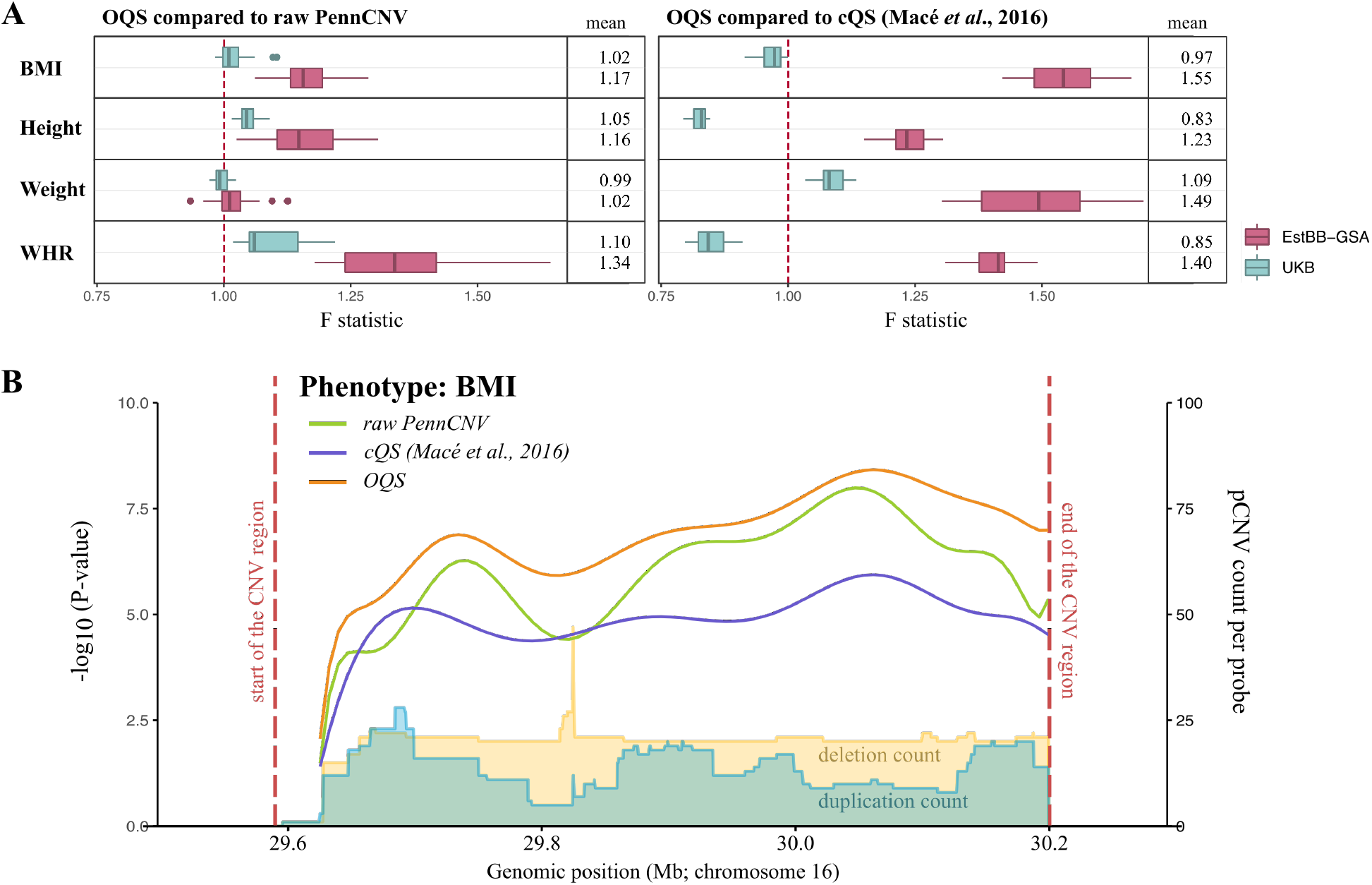
(A): The change of variance explained in mirror type model when using OQS over raw PennCNV or cQS^19^ in both EstBB-GSA and UKB depicted as distribution of F statistics calculated by randomising the probe pruning priority order 20 times (see Methods). Explained variance is increased when F>1 and decreased when F<1. (B) Locus plot of a CNV region in 16p11.2 BP4-BP5 (red dashed lines – chr16:29,590,000-30,200,000 in GRCh37) associated with BMI in EstBB-GSA dataset. The lines indicate the –log10 association P-values using mirror model with raw PennCNV calls (green), cQS (blue) and OQS (orange). The yellow and light blue areas illustrate the frequency of PennCNV deletion and duplication counts, respectively, across the region.

## Discussion

Genotyping microarray data are frequently used for CNV calling and analyses but up to 48% of the calls from commonly used software, such as PennCNV, are not supported by other omics measures and are, therefore, likely false positives. To counter this, a quality score based on results overlap between three detection software tools has been developed (cQS)^19^. We aimed at improving the discriminatory capacity of this score by devising an omics-informed CNV quality score – OQS – that incorporates independent omics-based sources of evidence to identify high quality PennCNV CNV calls (pCNVs).

Datasets included in the development of our OQS include gene expression levels from RNA sequencing (*GE* metric), summed methylated and unmethylated intensities at CpG sites (*MET* metric), as well as CNVs detected from WGS reads (*WGS* metric). Each of these three approaches yielded a quality metric between 0 and 1 for every pCNV, all of which showed high concordance. We found that the correlation between *WGS* and the other two metrics were ≥ 0.7, suggesting that the use of methylation and gene expression data for CNV quality assessment is a suitable alternative if WGS data are not available. Interestingly, the correlation between *WGS* and previously published cQS is quite low for comparison (0.17 for deletions, 0.43 for duplications) illustrating the potential benefit of incorporating omics data in CNV quality assessment compared to simple overlap between several detection software (which are all prone to same weaknesses, such as prioritisation of longer and discarding shorter CNV regions).

Using *GE, MET* and *WGS* metrics, we built predictive models relying only on output parameters of PennCNV that allow estimating CNV quality in datasets where no additional omics data are available. Although larger omics data sizes can lead to better CNV quality models, we believe that even modest sample sizes can be used in case the assessed set of CNVs are a good representative of the final CNV set in the analysis. In our study, the best quality models were built only on 441 pCNVs from LLDeep dataset having both *GE* and *MET* metrics calculated.

In validation sets of close relatives, OQS clearly discriminated between ‘familial’ (true positives) and ‘non-familial’ pCNVs, the former being attributed higher OQS compared to cQS. This effect was consistent over all independent tested datasets. Based on out-of-sample AUC and correlations, predicting quality of deletions was easier than predicting that of duplications. Possible explanations include better detection of deletions by PennCNV due to larger relative difference in allelic intensity ratio between one and two copies compared to two and three copies, or stronger effect of deletions on gene expression and methylation. In a second step, we compared OQS to raw PennCNV calls and cQS through an association analysis exercise aiming at replicating previously established CNV-trait associations. We found that OQS systematically increases (up to 34% in the EstBB-GSA and 10% in the UKB) the amount of explained variance when compared to raw PennCNV. Compared to cQS, we observed a strong improvement in explained variance in the EstBB-GSA (up to 55%) but not so in the UKB (down by 18%). As the associations we aimed at replicating were originally detected in the UKB using cQS approach, cQS-based associations suffer from Winner’s curse, which distorts the effect magnitudes in favour of the cQS. Alternatively, different quality scores might perform better in different datasets and combining the two might be a good option (e.g., by incorporating the maximum of the two scores in the analyses).

It is to be expected that the improvements offered by the OQS is small when studying strong associations in well-known and -detectable genomic region, as we have done. We expect to see greater improvement in intermediate-quality CNV regions for which previous studies have lacked statistical power for CNV-trait association detection. Additionally, as false pCNVs should be distributed randomly and uniformly across the genome, they only introduce a modest amount of noise per probe/region. However, given the difficulty to detect CNVs, which themselves tend to be rare, even slight improvement in statistical power to detect CNV associations can be beneficial.

Although we strived to optimise our models for different genotyping array types and densities, our results may still be specific to the arrays we explored. Furthermore, while the OQS helps to reduce false positive calls, it does not improve the false negative rate. Using only PennCNV output parameters as predictors is a limiting factor in itself as their prediction ability can vary from dataset to dataset. Furthermore, PennCNV detection algorithm considers each sample separately and does not exploit between-sample similarities which was shown to improve the detection of short (and frequent) CNV considerably^15^. Still, our omics-informed CNV quality assessment approach is not limited to PennCNV but can be used with any CNV detection method that produces multiple output parameters.

In conclusion, we developed a modular and customizable omics-based quality score framework that can be used for both genome-wide and smaller-scale CNV analyses. The OQS developed in the current study can be applied directly to filter out high confidence false positive pCNVs using a hard cut-off (i.e., OQS<0.5) or plugged into dosage-based association models, eliminating the need for an arbitrary CNV quality threshold. In turn, lower false negative rates increase statistical power to detect associations between CNVs and complex traits. Alternatively, with at least one suitable multi-omics measurements available for a subset of the samples, researchers can use our framework to build their own custom models, which could be applied to any CNV detection software, leading to further improved results for follow-up analyses.

## Supplementary Data

Supplementary data include eight notes, twelve tables and twelve figures.

## Consortia

### Estonian Biobank Research Team

Andres Metspalu, Tõnu Esko, Mari Nelis, Lili Milani

## Acknowledgements

We thank participants of Estonian Biobank (EstBB), UK Biobank (UKB), SkiPOGH and Lifelines Deep (LLDeep) for their data provision. The scholarships for PhD student mobility of M.L. and K.L. to work with SkiPOGH (University of Lausanne, Switzerland) and LLDeep (University of Groningen, The Netherlands) data, as well as the funding for EstBB data acquisition, was provided by the European Regional Development Fund (Project No. 2014-2020.4.01.16-0125 and Project No. 2014-2020.4.01.15-0012). This project has benefited from the funding of European Union’s Horizon 2020 Research and Innovation Programme under grant agreements No 101016775 and No 633666, and of the Estonian Research Council grant PUT (PRG687, PRG555 and PUT JD817). Z.K. was supported by the Department of Computational Biology at the University of Lausanne. M.N. was supported by Jacobs Foundation Research Fellowship (grant #2016 1217 09, to external collaborator Dr Katrin Männik, Health 2030 Genome Center, Geneva, Switzerland). The SKIPOGH study was supported by grants from the Swiss National Science Foundation (Grant Numbers 33CM30-124087 and 33CM30-140331). EstBB and part of the UKB computations were carried out on the High-Performance Computing (HPC) Center, University of Tartu. UKB association study was carried out on the JURA server, University of Lausanne. SkiPOGH computations were carried out on the HPC1 computational server of the University Hospital of Lausanne. We thank UMCG Genomics Coordination Center, the UG Center for Information Technology and their sponsors BBMRI-NL & TarGet for storage and compute infrastructure for LLDeep data. Finally, we thank Natàlia Pujol Gualdo and Triin Laisk for critical reading of the manuscript.

## Author Contributions

M.L., K.L., R.M. and Z.K. designed the study; M.L. conducted the main analyses including CNV detection, CNV quality assessment, modelling and association study in EstBB; C.A. performed the association study in UKB; Estonian Biobank Research Team coordinated the EstBB data generation and genotype data quality control; K.L. performed the quality control of EstBB gene expression; U.V., L.F. and C.W. contributed with LLDeep data acquisition, A.C. and K.L. performed the initial quality control; M.B. and S.S. generated and quality controlled SkiPOGH data; C.A., E.P and M.N. assisted with genotyping array-based CNV detection; M.K. performed CNV detection from whole genome sequencing reads; T.J., H.P., J.V., K.L. and Z.K. assisted with statistical analyses; M.L. drafted and C.A., A.P.M., U.V., R.M. and Z.K. made critical revisions to the manuscript; All authors read, approved, and provided feedback on the final manuscript.

## Data and Code Availability

Access to the UK Biobank Resource is available by application (http://www.ukbiobank.ac.uk/). Lifelines Deep RNA sequencing and methylation data can be accessed via European Genome-Phenome Archive (accession code EGAS00001001077). EstBB, SkiPOGH and LLDeep individual data are available upon request. The following publicly available software were used: PennCNV (http://penncnv.openbioinformatics.org/en/latest/), Genome Strip (https://software.broadinstitute.org/software/genomestrip/), QTLtools v1.1 (https://qtltools.github.io/qtltools/). Custom R code will be made available on GitHub.

