## Supplemental information for "Omics-informed CNV calls reduce false positive rate and improve power for CNV-trait associations"

### Supplementary information

### Supplementary Notes

#### S1. Estonian Biobank (EstBB): overview, CNV detection and quality control

*Dataset overview*

EstBB[1] is an Estonian population-based cohort of over 200,000 adult (aged ≥18 at recruitment) individuals recruited in years 2002-2020. All samples were genotyped using Illumina Global Screening Array (GSA) v1.0, GSA v2.0, GSA v2.0_ESTChip, and GSA v3.0_ESTChip2 arrays in 12 batches at the Core Genotyping Lab of the Institute of Genomics, University of Tartu. A subset of ~7,750 individuals recruited before 2012 have additionally been genotyped using Illumina Infinium OmniExpress-24 genotyping array. Another subset of ~2,500 individuals have whole-genome sequencing (WGS) data available[2]. The overlap between OmniExpress and WGS samples is ~1,000 individuals. Additionally, as part of smaller initiatives, RNA sequencing[3] and methylation (Infinium Human Methylation 450k Beadchip) data have been generated for subsets of the OmniExpress samples. Altogether, 1,066 OmniExpress samples have at least one out of WGS, methylation and RNA sequencing datasets available and are referred to as the EstBB multi-omics set or EstBB-MO.

*Quality control of GSA and OmniExpress CNV samples*

For both the OmniExpress and GSA datasets, we excluded samples with genotype call rate <98%, Hardy-Weinberg equilibrium test P-value $<1\times10^{-4}$ or mismatched sex based on chromosome X heterozygosity. Only one of each duplicated samples was retained. We created the intensity files (log R ratios (LRR) and B allele frequencies (BAF)) with Illumina GenomeStudio v2.0.4. For GSA samples, genotypes were re-clustered by manual realignment of cluster locations.

Autosomal copy number variations (CNVs) were called for 7,509 OmniExpress samples and 193,844 GSA samples using PennCNV detection software[4]. We denote the (true and false CNV) regions output by PennCNV as pCNV. The `hhall.hmm` (included in PennCNV software package) was chosen for the Hidden Markov Model (HMM) file and GC model file was created from `hg19.gc5Base.txt` file that was downloaded from the UCSC Genome Browser (https://hgdownload.soe.ucsc.edu). For OmniExpress, the population frequency of the B allele (PFB) file was created using all samples. Adjacent CNV calls were merged. We excluded all OmniExpress samples with >200 pCNV calls and with pCNV length >10Mb. For

GSA, the PFB file was created based on 1,000 randomly selected individuals from the first batch. Again, adjacent CNV calls were merged. Two full genotyping batches were found to be outliers based on the intensity signal compared to the rest of the batches and, thus, were excluded. Genotyping plates with >3 samples with either >200 called pCNVs or a total length of pCNV calls >10 Mb were excluded. Individual samples meeting these criteria were further removed. In total, 7,396 OmniExpress and 156,254 GSA samples were retained after filtering. The size of the sample overlap between the two datasets was 3,881 (out of which 39 samples belonged to the EstBB-MO subset).

*Relatives extraction and exclusion*

Relative pairs were detected using SNV genotypes with KING-robust kinship estimator[5] included in the PLINK 2.0 software (https://www.cog-genomics.org/plink/2.0/). We extracted monozygotic (MZ) twins (N=312 individuals; KING coefficient >0.354) and a subset of samples consisting of first-degree relatives (N=79,903 individuals; KING coefficient 0.177-0.354) from GSA samples. We excluded samples included in the EstBB-MO set from the OmniExpress samples and extracted the relative pairs with KING coefficient >0.177 (N=504 individuals). In order to create a GSA association study dataset, we excluded one sample of each pair with kinship coefficient >0.0884. This resulted in 89,516 unrelated GSA samples identical to what is used in[6].

*Phenotype data*

We extracted four anthropometric measurements (body mass index (BMI), height, weight and waist-to-hip ratio (WHR)) for each of the GSA association study samples. BMI, height and weight were collected during the biobank recruitment, while WHR was parsed from health registries and doctors' notes. If several WHR measurements were available, the most recent was retained. Phenotypes were inverse normal transformed and residualised on sex, age, age$^2$, genotyping batch and first 20 principal components.

#### S2. Lifelines Deep (LLDeep)

LLDeep is a subset of ~1,500 unrelated deeply phenotyped samples from the Dutch Lifelines cohort. All samples were genotyped using HumanCytoSNP-12 array and have RNA sequencing[7] and methylation (Infinium Human Methylation 450k Beadchip)[8] data available. The genotype dataset, as well as the quality control steps of omics data, are described elsewhere[9]. CNVs were detected in two batches of 865 and 522 samples. The `hhall.hmm` was chosen for the HMM file and GC correction was applied. The full dataset was included in the analyses as no individuals had >200 pCNVs or pCNVs longer than 10Mb.

#### S3. Swiss Kidney Project on Genes in Hypertension (SkiPOGH)

SkiPOGH is a family and population-based study of genetic determinants of blood pressure and renal function[10,11]. The samples were collected between 2009 and 2013 from the cantons of Bern and Geneva, and from the city of Lausanne. It contains 1,128 adult (aged ≥18) samples from 274 families (mostly trios) and is part of a larger European Project on Genes in Hypertension (EPOGH) study. All samples were genotyped using a dense Illumina 2.5 array. CNVs were detected in one batch of 675 samples using PennCNV software. The `hhall.hmm` was chosen for the HMM file and GC correction was applied. All carriers with >200 pCNVs or pCNVs longer than 10Mb were excluded resulting in 466 samples. Out of these, 405 had gene expression and 148 had methylation data available[12]. A separate set was compiled with all parent-child pairs that meet the CNV filtering criteria (319 samples from 102 families).

#### S4. UK Biobank (UKB): overview, CNV detection and filtering

*Dataset overview*

UKB is a population-based cohort of ~500,000 individuals from United Kingdom aged between 40 and 69 at recruitment. The majority of the samples (~450,000) are genotyped on Affymetrix UK Biobank Axiom array while the rest (~50,000) are genotyped on Affymetrix UK BiLEVE Axiom array. The dataset and general genotype quality control steps have been described by [13]. Although (exome) sequencing data is now available for a subset of UKB samples, this was not the case at the time of most of our analyses.

*CNV detection and quality control for familial analysis*

CNVs were detected in 106 batches using PennCNV. Individual specific intensity files containing B allele frequencies (BAF) and log R ratios (LRR) per probe were created using PennCNV-Affy conversion pipeline prior CNV detection. The PFB files were created for each batch separately, using 250 randomly selected samples per batch. The `hhall.hmm` was used and no GC correction was performed. Finally, adjacent CNV calls were merged. All samples with >200 pCNVs or pCNV total length >10Mb were excluded. Altogether, 401,571 samples were retained. Relative pairs were calculated using KING-robust kinship estimator[5].

Analogously to EstBB-GSA samples, we extracted MZ twins (N=302; coefficient >0.354) and first-degree relatives (N=42,032; KING coefficient 0.177-0.354).

*CNV detection and quality control for association analyses*

The subset of UKB samples and anthropometric phenotypes used for association analysis was prepared in a separate pipeline as described in [6]. Namely, the CNVs were called in 106 batches using Affymetrix genome-wide 6.0 array HMM file and GC correction. Samples genotyped on plates with mean pCNV count per sample >100 were excluded. Samples with >200 pCNVs or a single pCNV >10Mb were further removed. After these quality control steps, 331,522 unrelated British samples were retained. Four anthropometric traits were extracted and inverse normal transformed prior correction for sex, age, $age^2$, genotyping batch, and PC1-40.

#### S5. CNV detection from whole-genome sequencing reads

The Genome STRiP pipeline[14] was used to call CNVs in five separate batches for EstBB-MO samples. Eleven samples with excessive number of calls (#calls/#samples > median (across all samples) + 3 median absolute deviation) were removed. The union of the discovered sites was genotyped with Genome STRiP SVGenotyper module in all batches separately and merged. Duplicate calls were removed using the standard Genome STRiP duplicate removal settings, involving site overlap greater than 50% and duplicate score (logarithm of odds (LOD) score of genotype concordance at most discordant sample) greater than zero. In addition, low-quality (LQ) CNVs and CNVs with call rate less than 90% were excluded. Additionally, the pipeline excludes deletions shorter than 1,000bp and duplications shorter than 2,000bp.

#### S6. Preparations of methylation data

Methylation data from Infinium Human Methylation 450k Beadchip was collected for EstBB-MO, LLDeep and SkiPOGH samples. We extracted the methylation data matrices containing methylated and un-methylated intensities from sample-specific *idat* files using R package *minfi*[15] and summed those matrices to an overall intensity matrix. We eliminated all Type I methylation probes (N=135,476), since these cannot be used for capturing overall summed intensity, and only used Type II probes (N=350,036) in our study. All intensities that had detection P-values > $1\times10^{-16}$ were marked as missing and all probes with >5% missingness were filtered out. We corrected the overall intensity of each probe for age and sex. In EstBB-MO and SkiPOGH we additionally corrected for first four genotype principal components (PCs).

To exclude as much noise as possible while retaining the effect of CNVs on methylation data, we tested correcting the data for several numbers of methylation intensity PCs, ranging from 0 to 200 (using the following fixed categories: 0, 1, 2, 3, 4, 5, 10, 20, 30, 40, 50, 100, 150 and 200). CNV quality metrics based on methylation data (*MET* metric; as described in the main text) were calculated for each set of PCs. In each dataset, we chose the best set of PCs by maximising correlation with previously published consensus-based CNV quality score (cQS) [16] or *WGS* metric in EstBB-MO samples. The best number of PCs for *MET* metric was 30 for EstBB-MO deletions, 10 for EstBB-MO duplications, 100 for LLDeep deletions, 30 for LLDeep duplications, 1 for SkiPOGH deletions and 4 for SkiPOGH duplications (see **Figure S1**). Note that results obtained with different numbers of PCs did not vary substantially.

#### S7. Preparations of gene expression data

RNA-sequencing counts were obtained for EstBB-MO, LLDeep and SkiPOGH samples. We further processed the data as follows: (1) we removed genes with low or no expression in the majority of individuals by requiring for each gene to have $\geq 5$ individuals with a count per million (cpm) value greater than 0.5; (2) we normalized remaining genes by weighted trimmed mean of M-values; (3) we calculated the log(cpm) values of each gene, and Z-transformed these values; (4) we corrected the resulting gene expression measures for covariates: age, sex, blood component values, and batch. In EstBB-MO and SkiPOGH we additionally corrected for genotyping PC1-4.

Analogously to methylation data, we corrected the remaining residual expression of each gene for up to 200 expression PCs (calculated on the residuals) to further clean the signal. Additionally, we corrected expression residuals of each gene for the gene's independent SNP *cis*-eQTL effects (P-value < 0.05) in EstBB-MO and LLDeep. For EstBB-MO set the *cis*-eQTL analysis was run on SNPs within 500kbp proximity to the transcription start site using QTLtools v1.1[17]. For LLDeep we used the already published eQTLs[7]. In SkiPOGH we did not correct for eQTLs. We calculated the scores both before and after eQTL corrections and saw that correlations obtained after corrections were slightly higher. We also tested only including genes that showed R>0.1 correlation with copy number in [18].

Gene expression based CNV quality metric (*GE* metric) was calculated for each set of PCs and filters. We only considered genes that had at least 80% overlap with a CNV. The best set was chosen by maximising the correlation between *GE* metric and previously published cQS[16] or *WGS* metric in EstBB-MO samples. The number of selected PCs for *GE* metric was 30 for

EstBB-MO deletions, 4 for EstBB-MO duplications, 30 for LLDeep deletions, 50 for LLDeep duplications, 40 for SkiPOGH deletions and 30 for SkiPOGH duplications (**Figure S2**).

#### S8. CNV quality modelling

We used the stepwise regression with forward selection as an algorithm to pick the best parameters into the final omics-informed CNV quality score (OQS) model. Our first objective was to compile the initial parameter set. We did that using the PennCNV output, which contains a set of characteristics and parameters for each of the detected CNV regions. All parameters are described in **Table S1**. Altogether, we tested eight different models with slightly varying initial parameter sets. First two initial parameter sets we tested included all PennCNV output variables as was done before[16]. The remaining initial sets were compiled using subsets of PennCNV output variables and additional variables calculated based on PennCNV output. Initial sets per model are shown in **Table S2**.

During each model-building step of the stepwise forward selection process, the following parameters remaining in the initial parameter set were tested:

a. PennCNV output (or derived) parameters not yet in the model;
b. interactions between a CNV-specific and a sample-specific parameter already in the model;
c. (if allowed) higher order terms of parameters already in the model.

An extra term was added to the model if it minimized the average mean square error (MSE) from *k*-fold cross-validation. If no term minimized the MSE, the algorithm stopped and returned the existing model. Models were built separately for deletions and duplications.

### Supplementary Tables

**Table S1**. Parameters used for CNV quality modelling. BAF – B allele frequency, LRR – log R ratio.

| **Parameter** | **Sample/variant-specific** | **Obtained** | **Variable name** |
|---|---|---|---|
| mean BAF | sample | from PennCNV output | `BAF_mean` |
| standard deviation of BAF | ” | ” | `BAF_SD` |
| BAF drift | ” | ” | `BAF_drift` |
| mean LRR | ” | ” | `LRR_mean` |
| standard deviation of LRR | ” | ” | `LRR_SD` |
| absolute waviness factor | ” | ” | `WF` |
| number of variant calls | ” | ” | `NumCNV` |
| variant confidence | variant | ” | `Max_Log_BF` |
| variant length | ” | ” | `Length_bp` |
| number of probes | ” | ” | `No_probes` |
| variant length per probe | ” | calculated as `Length_bp / No_probes` | `Length_per_Probe` |
| number of variant calls corrected for genotyping array size | sample | calculated as `NumCNV` / number of probes on array since number of variants per sample directly depends on array density | `NumCNV_corr` |
| corrected and dichotomised number of variant calls | sample | We noticed in several independent datasets that the CNV quality starts to decline rapidly for samples with number of CNVs (corrected for array density) over a certain threshold. We used ROC curves in order to pinpoint this threshold and found that in case of deletions, its value falls into a narrow range of $3.8\times10^{-5}...3.9\times10^{-5}$ dependent on the dataset (**Figure S4**). Therefore, we compiled a parameter for deletions that equals zero if `NumCNV_corr` is below and one if it is above the given threshold. | `NumCNV_bin` |

**Table S2**. Eight initial sets of explanatory variables included in the step-wise model selection algorithm. The explanatory variables are either output parameters of PennCNV software or directly calculated from these output parameters (**Table S1**). Two out of eight models allow higher order terms. All models allow interactions between sample-specific and CNV-specific parameters. Model building is described in **Supplementary Note S8**. The final set of parameters selected to the model depends on the dataset.

| Model No. | Initial parameter set | Higher terms |
|---|---|---|
| 1 | Sample-specific parameters, CNV confidence score, NumCNV, Length_bp, No_Probes | No |
| 2 | Sample-specific parameters, CNV confidence score, NumCNV, Length_bp, No_Probes | Yes |
| 3 | Sample-specific parameters, CNV confidence score, NumCNV, Length_per_Probe | Yes |
| 4 | Sample-specific parameters, CNV confidence score, NumCNV | No |
| 5 | Sample-specific parameters, CNV confidence score, NumCNV_corr, Length_bp, No_Probes | No |
| 6 | Sample-specific parameters, CNV confidence score, NumCNV_bin, Length_bp, No_Probes | No |
| 7 | Sample-specific parameters, CNV confidence score, NumCNV_bin, Length_per_Probe | No |
| 8 | Sample-specific parameters, CNV confidence score, NumCNV_bin | No |

**Table S3**. 21 CNV region–phenotype pairs that are analysed in the current study in the EstBB-GSA and UKB cohorts. All pairs reached an association P-value < $1\times10^{-4}$ in previously published study[19], using cQS[16], in at least one analysis type (mirror/ deletion only/ duplication only; flagged with "+"). In the EstBB-GSA and UKB columns, the "+" sign indicates that the corresponding association was also significant (for at least one probe in the CNV region) in the respective analysis of our current study. In this case the significance threshold was set to P-value < $0.05/21 = 2.38\times10^{-3}$ (using raw PennCNV calls). *In the EstBB-GSA, the CNV region on chromosome 18 contained a probe with P < 2.38e-3 but the CNV structure in the region and the association with BMI/weight was clearly not the same as reported before [19]. Therefore, this region was not included in the EstBB-GSA follow-up analyses (**Figure S5**).

| **Phenotype** | **CNV region (hg37)** | **Macé et al. (2017)** | | | **EstBB-GSA** | | | **UKB** | | |
|---|---|---|---|---|---|---|---|---|---|---|
| | | **Deletions** | **Mirror** | **Duplications** | **Deletions** | **Mirror** | **Duplications** | **Deletions** | **Mirror** | **Duplications** |
| BMI | 16:28820000-29040000 | + | + | | | + | | + | + | |
| | 16:29590000-30200000 | + | + | | + | + | | + | + | |
| | 18:57660000-57900000 | + | + | | +* | +* | | + | + | |
| | 22:19030000-20310000 | | + | + | | | + | | + | + |
| | 22:20740000-21000000 | | + | + | | | | | + | + |
| Weight | 1:146530000-147430000 | + | + | | | | | + | + | |
| | 2:111580000-111780000 | | + | | | | | | + | |
| | 2:112680000-112780000 | | + | | | | | | + | |
| | 3:196220000-196420000 | | + | | | | | | | |
| | 16:28820000-29040000 | + | + | | | + | | + | + | |
| | 16:29590000-30200000 | + | + | | + | + | | + | + | |
| | 18:57660000-57900000 | + | + | | +* | +* | | + | + | |
| | 22:20740000-21000000 | | + | + | | + | | | + | + |
| Height | 1:146530000-147430000 | + | + | | + | + | | + | + | |
| | 3:196220000-196420000 | | + | | | | | | + | |
| | 3:196710000-196920000 | | + | | | | | | + | |
| | 11:27010000-27230000 | + | + | | + | + | | | | |
| | 15:22780000-23070000 | + | + | | | | | + | + | |
| | 16:29590000-30200000 | + | + | | + | + | | + | + | |
| WHR | 7:73020000-73110000 | | + | + | | | | | | |
| | 16:29590000-30200000 | + | + | | + | + | | + | + | |

**Table S4.** The number and percentage of PennCNV calls (pCNVs) that could be evaluated using methylation (*MET*), gene expression (*GE*) and whole-genome sequencing (*WGS*) based quality metrics in EstBB-MO, LLDeep and SkiPOGH datasets. The combined metric (*EXTR*) requires the availability of at least two out of three omics based metrics. The percentage is calculated as the fraction of pCNVs evaluated amongst all autosomal pCNVs of the set of individuals with respective omics dataset available (after sample exclusion in quality control). In case of *GE* metric, 80% gene-pCNV overlap is required.

| | **EstBB-MO** | **LLDeep** | **SkiPOGH** |
|---|---:|---:|---:|
| ***MET* metric** | 3,228 (41.8%) | 3,476 (68.7%) | 5,048 (44.9%) |
| ***GE* metric** | 462 (4.7%) | 706 (7.1%) | 1,497 (3.8%) |
| ***WGS* metric** | 23,977 (100%) | - | - |
| **Combined metric (*EXTR*)** | 3,496 | 441 | 410 |

**Table S5**. Number of pCNVs per combined (*EXTR*) metric range in three datasets.

| **Metric range** | **EstBB-MO** | | **LLDeep** | | **SkiPOGH** | |
|---|---|---|---|---|---|---|
| | **Deletions** | **Duplications** | **Deletions** | **Duplications** | **Deletions** | **Duplications** |
| [0;0.1] | 828 | 829 | 110 | 63 | 163 | 107 |
| (0.1;0.2] | 36 | 33 | 9 | 2 | 13 | 8 |
| (0.2;0.3] | 10 | 15 | 1 | 2 | 1 | 6 |
| (0.3;0.4] | 7 | 19 | 0 | 0 | 0 | 3 |
| (0.4;0.5] | 2 | 6 | 0 | 1 | 2 | 0 |
| (0.5;0.6] | 2 | 9 | 1 | 0 | 0 | 0 |
| (0.6;0.7] | 23 | 33 | 0 | 2 | 0 | 3 |
| (0.7;0.8] | 45 | 77 | 3 | 6 | 1 | 5 |
| (0.8;0.9] | 71 | 129 | 2 | 20 | 0 | 11 |
| (0.9;1.0] | 726 | 596 | 92 | 127 | 50 | 37 |

**Table S6**. Mean scores for familial and non-familial pCNVs calculated using models built on Estonian OmniExpress samples. The models are tested on monozygotic twins of UK Biobank and EstBB-GSA samples, as well as parent-child pairs form the SkiPOGH dataset. Altogether we tested eight models (**Table S2**) for three omics-based metrics (*WGS* – whole-genome sequencing metric, *MET* – methylation metric, *GE* – gene expression metric) and a combined metric (*EXTR*). The best model was selected as the one that maximises the difference between familial and non-familial pCNVs over three independent datasets. (Table in a separate .xlsx file)

**Table S7**. Mean scores for familial and non-familial pCNVs calculated using models built on LLDeep samples. Models are tested on monozygotic twins of UK Biobank and EstBB-GSA samples, as well as parent-child pairs form SkiPOGH dataset. Altogether we tested eight models (**Table S2**) for two omics-based metrics (*MET* – methylation metric, *GE* – gene expression metric) and a combined metric (*EXTR*). The best model was selected as the one that maximises the difference between familial and non-familial pCNVs over three independent datasets. The best model is highlighed in blue. (Table in a separate .xlsx file)

**Table S8**. Description of best omics-informed CNV quality model for deletions. All parameters are explained in **Table S1**. 'X:Y' denotes the interaction term between parameters X and Y.

| Parameter | Effect size (beta) |
|---|---|
| (Intercept) | -2.12350638759242 |
| Max_Log_BF | 0.138074328580999 |
| LRR_mean | 192.33129681258 |
| NumCNV_bin | -1.17079719833943 |
| Max_Log_BF:LRR_mean | -6.07975553625751 |

**Table S9**. Description of best omics-informed CNV quality model for duplications. All parameters are explained in **Table S1**. 'X:Y' denotes the interaction term between parameters X and Y.

| Parameter | Effect size (beta) |
|---|---|
| (Intercept) | -114.039850181999 |
| Max_Log_BF | -0.0368788122813245 |
| Length_bp | -9.76978925613445e-06 |
| LRR_SD | -34.7848931008816 |
| BAF_mean | 233.943409217459 |
| BAF_drift | -2032.95959074771 |
| Max_Log_BF:LRR_SD | 0.60672686014698 |
| Length_bp:LRR_SD | 0.000111986438009925 |

**Table S10**. CNV associations analysis results for three settings (raw PennCNV, cQS[16] and omics-informed quality score (OQS)) in EstBB-GSA dataset. 21 CNV region-phenotype pairs ($P<1\times10^{-4}$; [19]) are included in the analysis (**Table S3**). (Table in a separate .xlsx file)

**Table S11**. CNV associations analysis results for three settings (raw PennCNV, cQS[16] and omics-informed quality score (OQS)) in UKB dataset. 21 CNV region-phenotype pairs ($P<1\times10^{-4}$; [19]) are included in the analysis (**Table S3**). (Table in a separate .xlsx file)

**Table S12**. F statistics that characterise the change in explained variance when comparing OQS to raw PennCNV and cQS[16] in EstBB-GSA and UKB datasets. F>1 means an increase and F<1 a decrease in explained variance (i.e. F=1.16 indicates a 16% increase in explained variance). The values presented in this table are calculated as an average over 20 runs. None of the values reach even nominal statistically significance (P<0.05) due to the low number of independent associated regions per phenotype.

| Phenotype | Analysis type | EstBB-GSA | | UKB | |
|---|---|---|---|---|---|
| | | *F (OQS vs raw PennCNV)* | *F (OQS vs cQS)* | *F (OQS vs raw PennCNV)* | *F (OQS vs cQS)* |
| BMI | Mirror | 1.17 | 1.55 | 1.02 | 0.97 |
| Height | Mirror | 1.16 | 1.23 | 1.05 | 0.83 |
| Weight | Mirror | 1.02 | 1.49 | 0.99 | 1.09 |
| WHR | Mirror | 1.34 | 1.40 | 1.10 | 0.85 |
| BMI | Deletion only | 1.19 | 1.30 | 1.02 | 0.93 |
| Height | Deletion only | 1.08 | 1.09 | 0.97 | 0.96 |
| Weight | Deletion only | 1.33 | 1.46 | 1.09 | 0.88 |
| WHR | Deletion only | 1.13 | 1.23 | 1.10 | 0.82 |
| BMI | Duplication only | 0.85 | 1.43 | 1.00 | 1.01 |
| Weight | Duplication only | - | - | 1.03 | 1.02 |

### Supplementary Figures

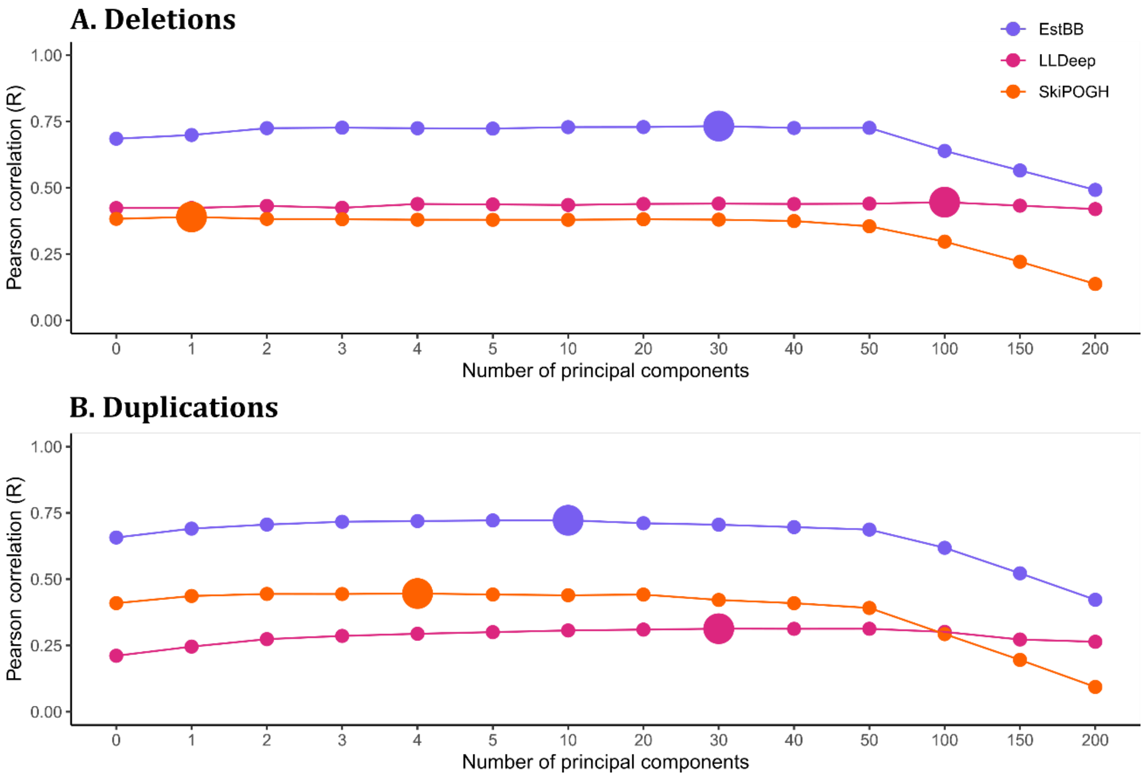

**Figure S1**. Methylation metrics for (A) deletion and (B) duplication quality (*MET*) calculated using 0 to 200 methylation intensity principal components (PCs; **Supplementary Note S6**) and correlated (Pearson's correlation) with *WGS* metric in EstBB-MO and cQS [16] in LLDeep and SkiPOGH. The number of PCs chosen for further analyses are indicated with large points. We used *MET* values corrected for 30 PCs for EstBB-MO deletions, 10 PCs for EstBB-MO duplications, 100 PCs for LLDeep deletions, 30 PCs for LLDeep duplications, 1 PCs for SkiPOGH deletions and 4 PCs for SkiPOGH duplications, as these maximised the corresponding correlations in the respective datasets.

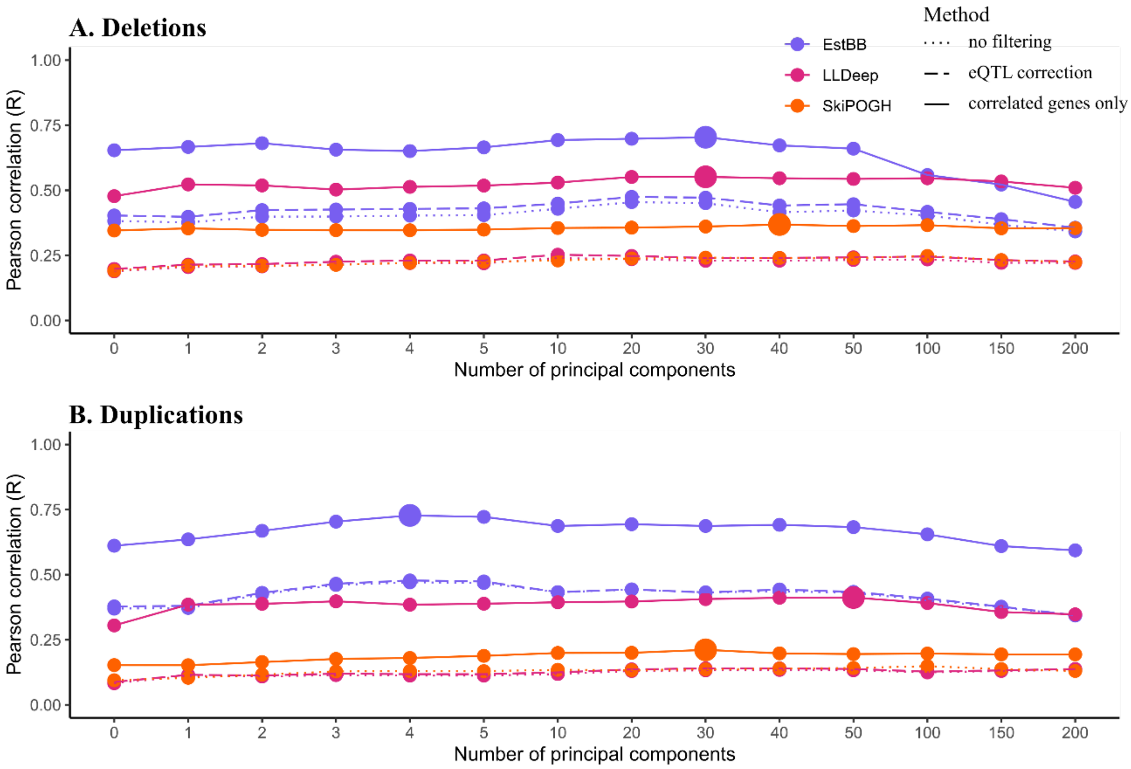

**Figure S2**. Gene expression metrics for (A) deletion and (B) duplication quality (*GE*) calculated using 0 to 200 expression principal components (PCs; dotted line). Additionally we tested correcting gene expression for eQTLs prior score calculations (dashed line) and only including genes which showed correlation to CNV status in [18] (solid line; **Supplementary Note S7**). We correlated (Pearson's correlation) *GE* values with *WGS* metric in EstBB-MO and cQS [16] in LLDeep and SkiPOGH. In most cases, correcting for eQTLs slightly increased the correlation while gene filtering improved the correlations significantly. The number of PCs chosen for further analyses are indicated with large points. We used *GE* metric values corrected for eQTLs and 30 PCs for EstBB-MO deletions, 4 PCs for EstBB-MO duplications, 30 PCs for LLDeep deletions, 50 PCs for LLDeep duplications, 40 PCs for SkiPOGH deletions and 30 PCs for SkiPOGH duplications, as these maximised the corresponding correlations in the respective datasets.

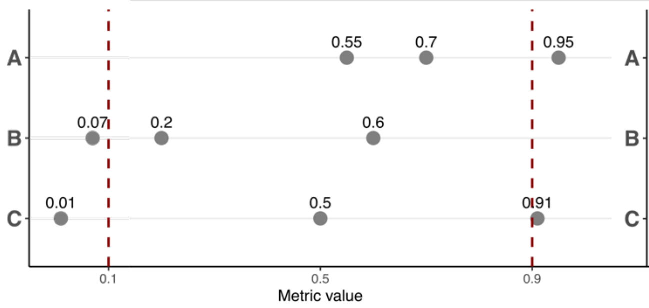

| | MEAN | MAX | EXTR | EstBB-MO | LLDeep |
|---|---|---|---|---|---|
| A | 0.73 | 0.95 | 0.95 | 18.2% | 11.3% |
| B | 0.29 | 0.60 | 0.07 | 15.2% | 18.4% |
| C | 0.47 | 0.91 | 0.01 | 2.2% | 2.9% |

**Figure S3**. For each PennCNV call (pCNV), we calculated up to three omics-based CNV quality metrics (*GE*, *MET* and *WGS*). Our aim was to combine them into one metric per pCNV. We considered using the mean (*MEAN*), maximum (*MAX*) and the most extreme (i.e., furthest from 0.5, *EXTR*) metric. We divided the metric value range into three subranges: false positive [0; 0.1), ambiguous [0.1; 0.9] and true positive (0.9, 1]. For the majority of pCNVs (64.4% in EstBB-MO (full set N=3,496) and 67.3% in LLDeep (N=441)) all omics-based metrics were in close agreement with all values falling into the same subrange. For these cases the exact choice of combined metric makes little difference. Here we bring three toy examples of cases where the metrics do not agree: (A) at least one metric indicates a true positive CNV while others are ambiguous; (B) at least one metric indicates a false positive CNV while others are ambiguous; (C) at least one metric indicates a false positive CNV and at least one other a true positive CNV. Each example is followed by the values of corresponding combined metrics and the percentages on pCNVs that fall under these example categories in EstBB-MO and LLDeep. The usage of *MEAN* metric dilutes the effect of omics-based metrics that clearly indicate the presence of a true or false positive CNV, which is only reasonable in example C (2.2%-2.9% of cases). The usage of *MAX* metric is only intuitive when we expect the CNV to be true positive (example A; 11.3%-18.2% of cases). The usage of *EXTR* metric is intuitive in both examples A and B (29.7%-33.4% of cases) and even in example C it produces correct estimations in roughly half of the cases. Therefore, we chose to use *EXTR* as our combined metric per pCNV.

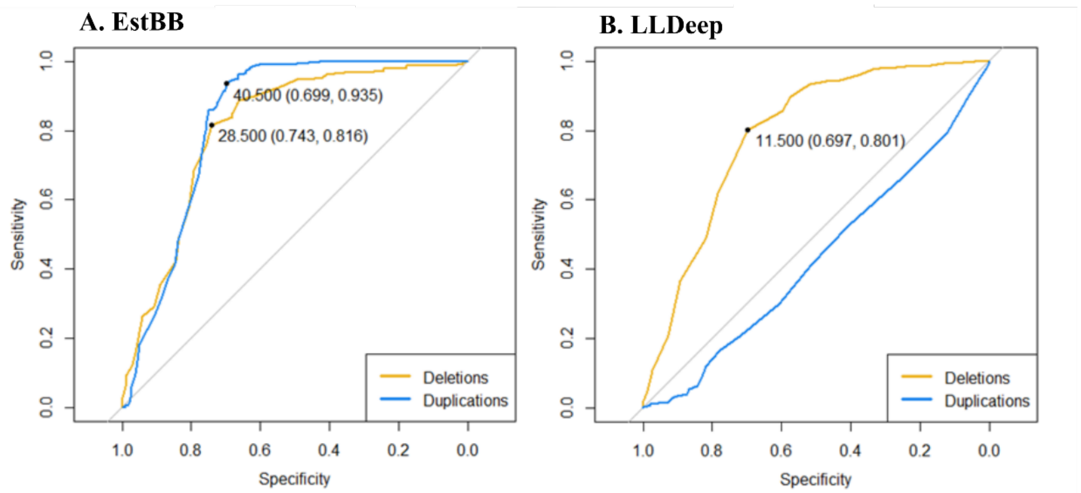

**Figure S4**. ROC curves built on (A) EstBB-MO and (B) LLDeep pCNVs with number of pCNVs per person as predictor. The true and false calls are defined as methylation metric *MET* >0.9 and <0.1, respectively. We used *MET* metric as it was present in both datasets in greater extent that combined metrics (the results for combined metric did not differ significantly). For duplications (blue) in EstBB-MO and deletions (yellow) in both datasets, there is a clear cut-off for the number of pCNVs per sample that can be used to discriminate between true and false calls. These cut-offs values are presented as dots on the ROC curves (followed by the corresponding specificity and sensitivity values). The exact cut-off is not generalisable as it is dependent on array density (~730k for EstBB-MO and ~300k for LLDeep). After correction for array density, the values for deletions are comparable: $3.9\times10^{-5}$ in EstBB and $3.8\times10^{-5}$ in LLDeep. We used these values to create a binary variable out of number of CNVs corrected for array density.

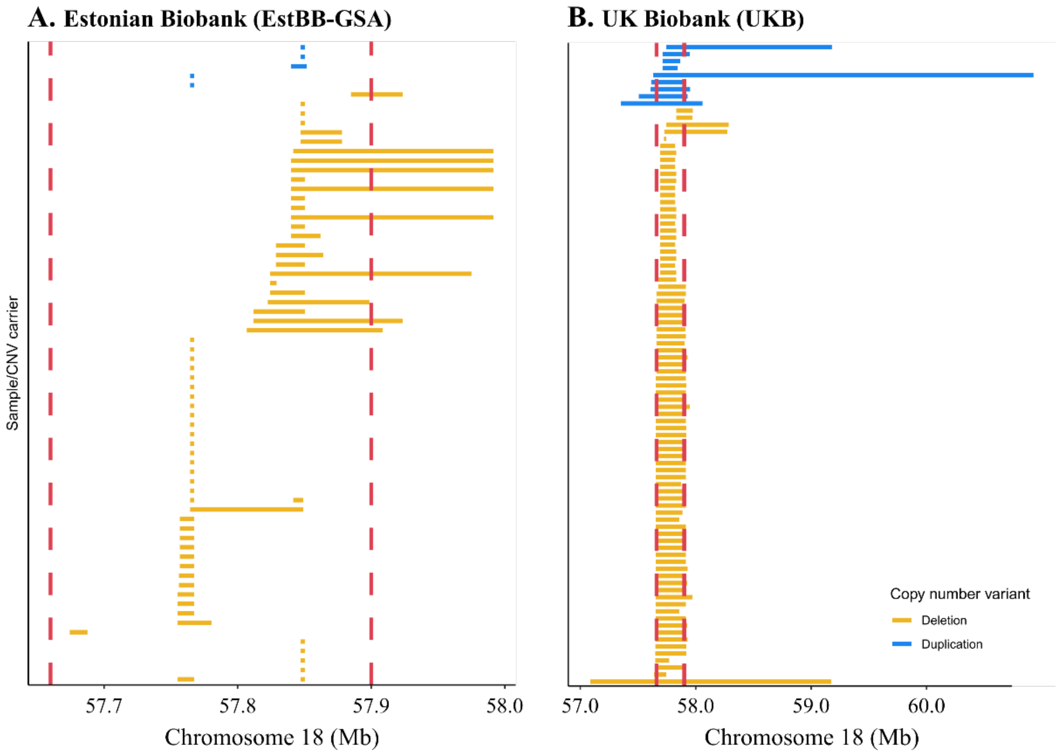

**Figure S5**. In EstBB-GSA analyses we excluded the 18q21.32 CNV previously associated with body mass index and weight[19]. Although there were EstBB-GSA CNVs (A) overlapping the region of interest (indicated with red dashed lines), they are not the CNVs for which the association was detected. In UKB (B) this region was included in the association tests.

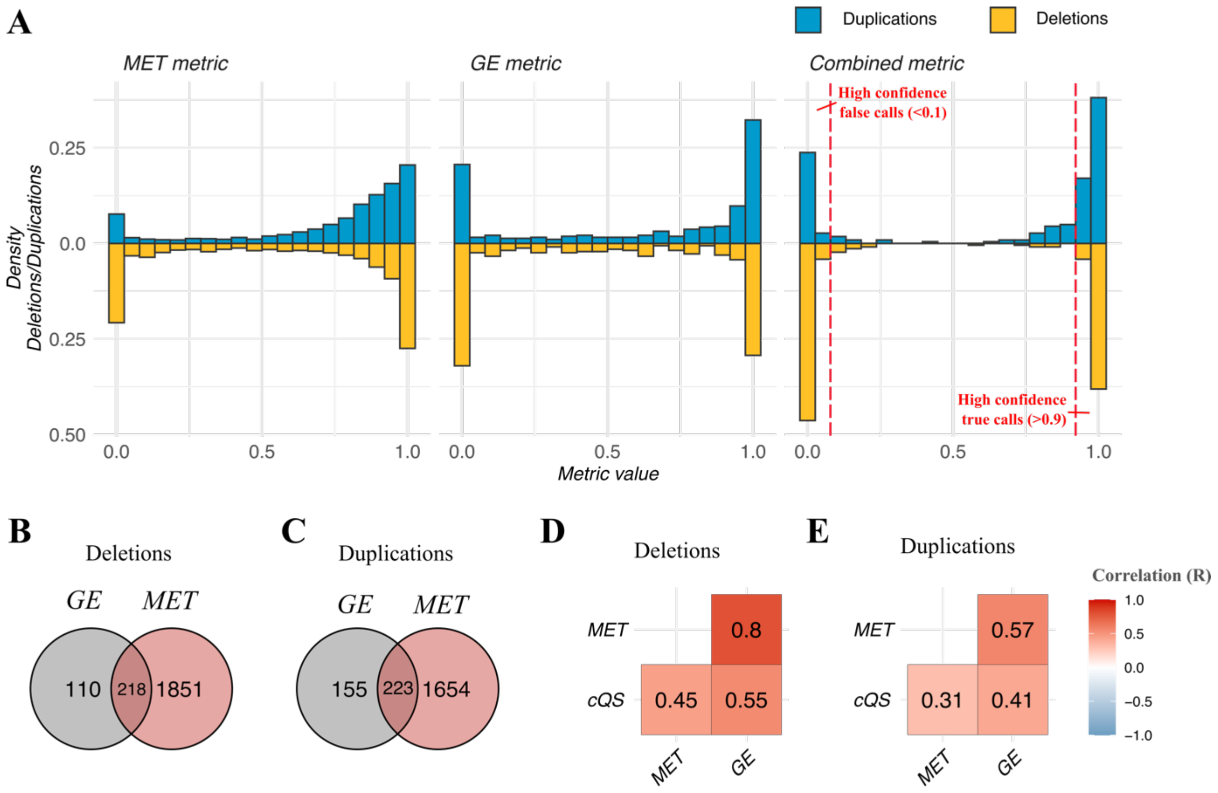

**Figure S6**. (A) Distributions of CNV quality metrics based on methylation (*MET*) and gene expression (*GE*) as well as their combination metric in LLDeep cohort for duplications (blue) and deletions (yellow). Altogether, 4,211 pCNVs have either *MET* or *GE* metric calculated. The combined metric is calculated for 441 pCNVs. 50.5% of deletions and 28.3% of duplications are high confidence false calls based on the combined metric (value <0.1), 42.2% of deletions and 57.0% of duplications are high confidence true calls (value >0.9). Number of deletions (B) and duplications (C) in LLDeep dataset that could be evaluated with *MET* and/or *GE* metrics. Pearson correlations between different quality metrics for deletions (D) and duplications (E), including with previously published cQS[16].

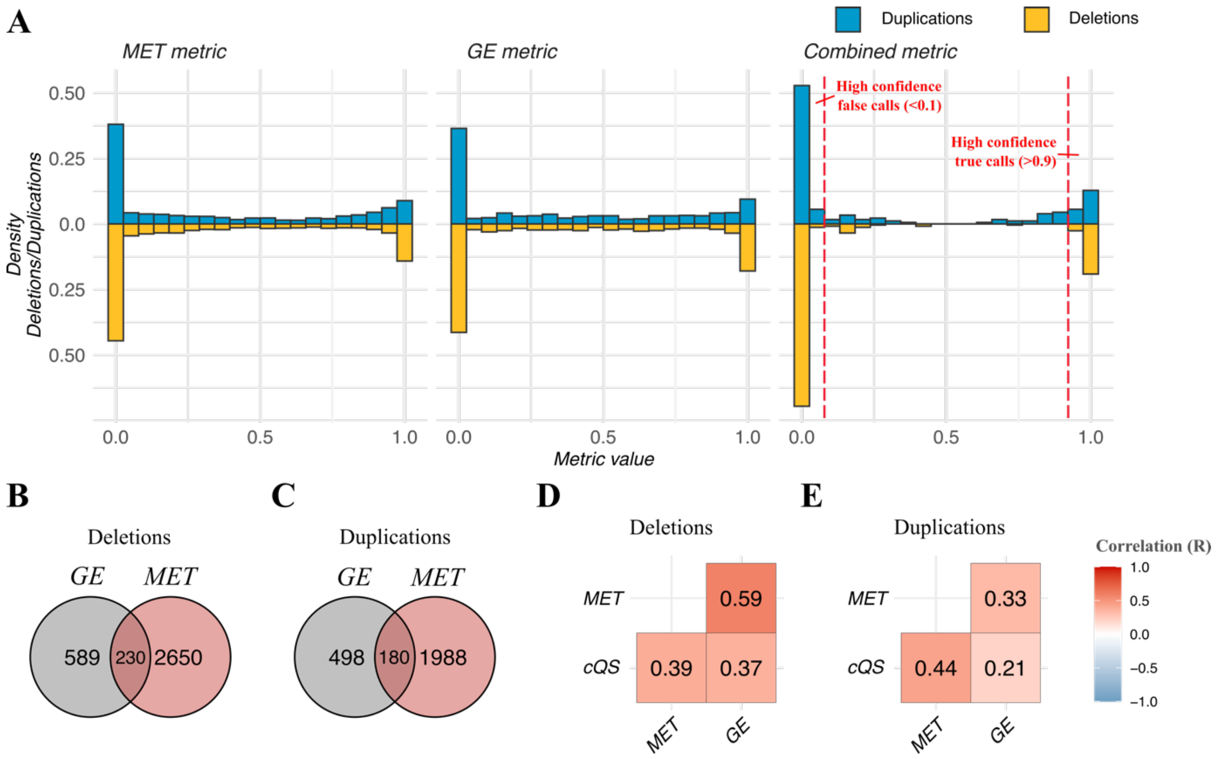

**Figure S7**. (A) Distributions of CNV quality metrics based on methylation (*MET*) and gene expression (*GE*) as well as their combination metric in SkiPOGH cohort for duplications (blue) and deletions (yellow). Altogether, 6,135 pCNVs have either *MET* or *GE* metric calculated. Combined score was calculated for 410 pCNVs. 70.9% of deletions and 59.4% of duplications are high confidence false calls based on at least one omics metric (value <0.1), only 21.7% of deletions and 20.6% of duplications are high confidence true calls (value >0.9). Since the fraction of false positive calls is much higher and true positive calls much lower in this dataset compared to others, we did not exclude SkiPOGH in CNV quality model building step. Number of deletions (B**)** and duplications (C) in SkiPOGH dataset that could be evaluated with *MET* and/or *GE* metrics. Pearson correlations between different quality metrics for deletions (D) and duplications (E), including with previously published cQS[16].

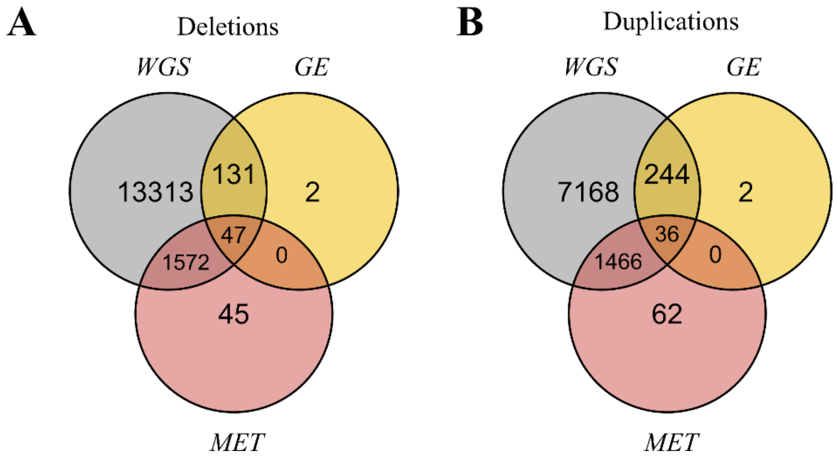

**Figure S8**. Venn diagrams with number of deletions (A) and duplication (B) that can be evaluated using whole-genome sequencing (*WGS*), methylation (*MET*) and/or gene expression (*GE*) metrics in EstBB-MO dataset. The combined metric (*EXTR*) is calculated for pCNVs that have at least two omics metrics available.

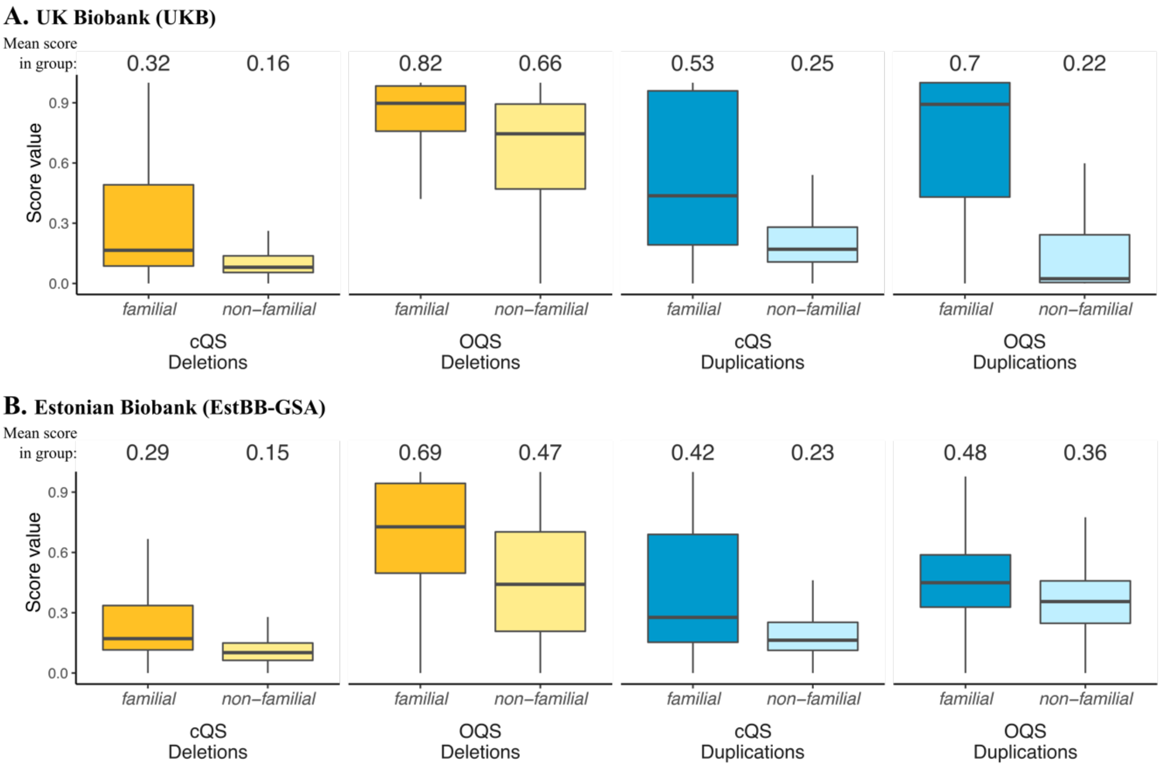

**Figure S9**. cQS[16] and omics-informed (OQS) CNV quality score comparisons between familial and non-familial pCNVs in (A) the UKB and (B) the EstBB-GSA first-degree relatives. Having a pCNV replicate in a family member is a good proxy for a pCNV to be a true call, while non-familial pCNV sets contain both true and false calls. The mean score of each pCNV group is shown on top of the figure. For both deletions (yellow) and duplications (blue), OQS shows higher scores for familial pCNVs compared to cQS. Outliers are not shown on the figure but are still included in the mean calculations.

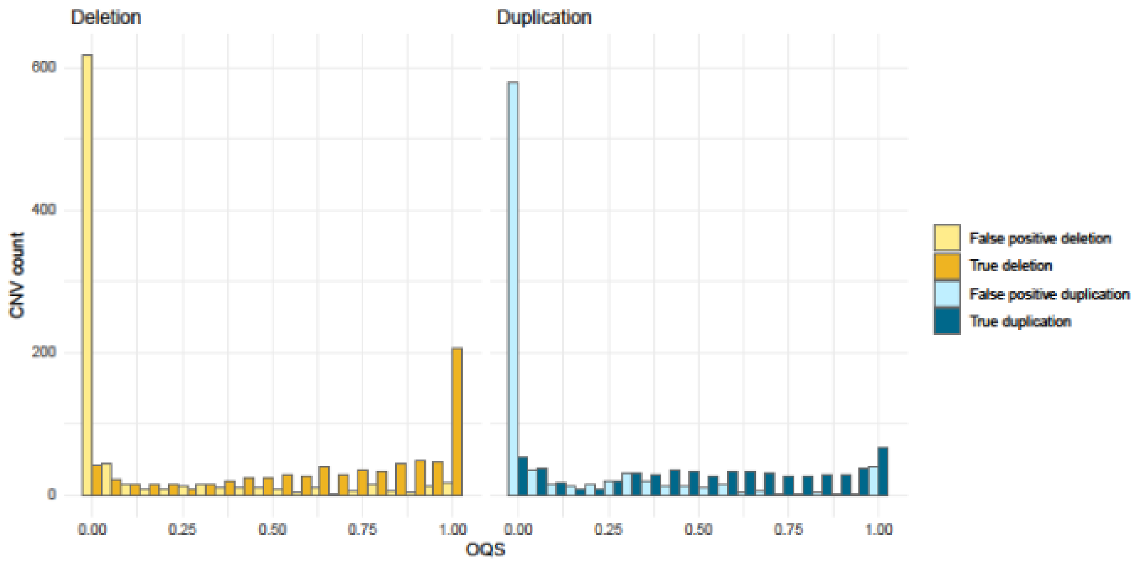

**Figure S10**. Distribution of predicted OQS in EstBB-MO dataset. The pCNV are divided into false positives (calculated omics-based *EXTR* metric < 0.1) and true CNVs (*EXTR* > 0.9).

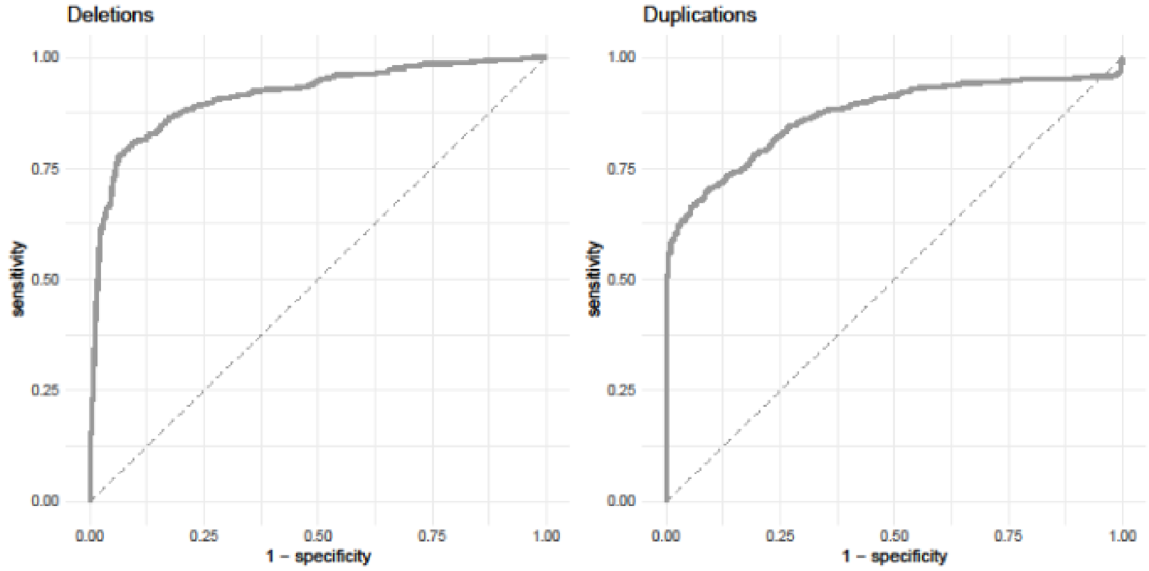

**Figure S11**. ROC curves of OQS in EstBB-MO dataset. The pCNV are divided into false positives (calculated omics-based *EXTR* metric < 0.1) and true CNV (*EXTR* > 0.9). For deletions AUC=0.91, for duplications AUC=0.87.

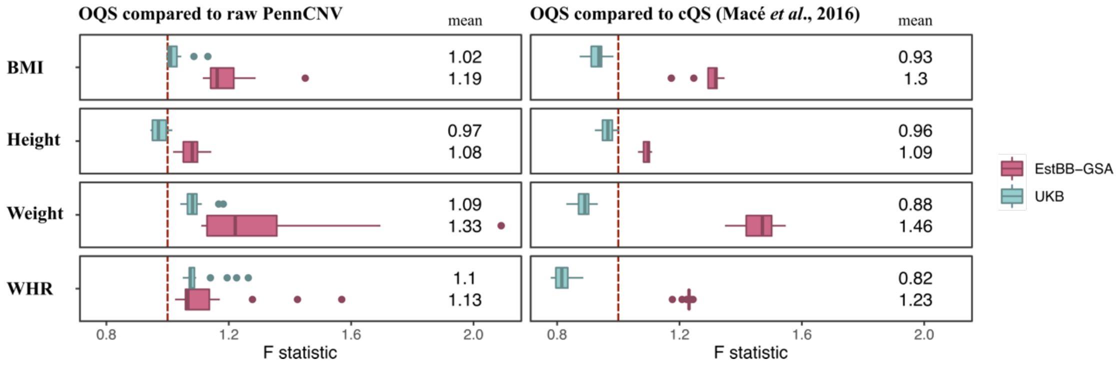

**Figure S12**. The change of variance explained in deletion-only model when using OQS over raw PennCNV or cQS[16] in both EstBB-GSA and UKB depicted as distribution of F statistics calculated by randomising the probe pruning priority order 20 times (see **Methods**). Explained variance is increased when F>1 and decreased when F<1.
